# Mutational analysis of the F plasmid partitioning protein ParA reveals novel residues required for oligomerisation and plasmid maintenance

**DOI:** 10.1101/2024.03.17.585406

**Authors:** Nivedita Mitra, Dipika Mishra, Irene Aniyan Puthethu, Ramanujam Srinivasan

## Abstract

Mobile genetic elements such as plasmids play a crucial role in the emergence of antimicrobial resistance. Hence, plasmid maintenance proteins like ParA of the Walker A type cytoskeletal ATPases/ ParA superfamily are potential targets for novel antibiotics. Plasmid partitioning by ParA relies upon ATP-dependent dimerisation and formation of chemophoretic gradients of ParA-ATP on bacterial nucleoids. Though polymerisation of ParA has been reported in many instances, the need for polymerisation in plasmid maintenance remains unclear. In this study, we provide novel insights into the polymerisation of ParA and the effect of polymerisation on plasmid maintenance. We first characterise two mutations, Q351H and W362E, in ParA from F plasmid (ParA_F_) that form cytoplasmic filaments independent of the ParBS_F_ partitioning complex. Both mutants fail to partition plasmids, do not bind non-specific DNA and act as super-repressors to suppress transcription from the ParA promoter. Further, we show that the polymerisation of ParA_F_ requires the conformational switch to the ParA-ATP* state. We identify two mutations, R320A in the C-terminal helix-14 and E375A helix-16 of ParA_F,_ that abolish filament assembly and affect plasmid partitioning. Our results thus suggest a role for higher-order structures or polymerisation of ParA in plasmid maintenance.

## INTRODUCTION

Antibiotic resistance is one of the major concerns for the current state of the world health system. The World Health Organization has declared antimicrobial resistance a global priority and identified Enterobacteriaceae as one of the major groups of pathogens as agents against which new antibiotics are urgently needed (Simonsen 2018; World Health Organization 2021). Enterobacteriaceae family includes *Klebsiella* and *Escherichia coli*, which are often acquired as hospital-borne infections and fail to respond to antibiotics due to the presence of multidrug resistance genes (Machado et al. 2022; Cerceo et al. 2016; Jalil and Al Atbee 2022). These genes are usually carried on single or low-copy number plasmids and are thought to be the most prevalent means to transfer antibiotic resistance genes across bacterial species (Yong et al. 2009; Phan et al. 2015; Stephens et al. 2020; Meynell et al. 1968; Moran et al. 2015). Thus, understanding plasmid maintenance and spread could be crucial to curtailing the rapid growth of multidrug resistance in bacteria.

Stable maintenance of these low copy number plasmids depends upon the functions of cytoskeletal proteins such as the actin homolog ParM (Møller-Jensen et al. 2003; van den Ent et al. 2002; Salje et al. 2010), tubulin homolog TubZ (Tinsley and Khan 2006; Anand et al. 2008; Larsen et al. 2007) and the Walker A type Cytoskeletal ATPases (WACA or the ParA superfamily)(Walker et al. 1982; Motallebi-Veshareh et al. 1990; Koonin 1993). The ParA family of proteins function to equipartition genetic material, both plasmids and chromosomes, during the bacterial cell cycle program (Ebersbach and Gerdes 2001; Castaing et al. 2008; Le Gall et al. 2016; McLeod et al. 2017; Lutkenhaus 2012; Lee and Grossman 2006). The F plasmid is one of the well-studied model systems to understand the mechanisms by which the ParA family of proteins function to ensure equipartitioning of the replicated plasmids into the daughter cells during cell division. ParA_F_, also known as SopA, is a member of the ParA superfamily, which binds to the nucleotide ATP and localises to the nucleoid within the bacterial cell (Hatano et al. 2007; Castaing et al. 2008; Roberts et al. 2012; Le Gall et al. 2016; McLeod et al. 2017; Hirano et al. 1998). ParA_F_ has an intrinsic ATP hydrolysis activity, which is stimulated several folds upon interaction with the partitioning complex comprising the adapter protein ParB_F_ (SopB) and the *parS_F_* (*sopC*) DNA sequence ^(^Ravin et al. 2003; Watanabe et al. 1992; Libante et al. 2001; Ah-Seng et al. 2013; Ogur^a^ and Hiraga 1983; Lane et al. 1987; Bouet et al. 2007; Barillà et al. 2005, 2007). Further, upon interaction with the SopBC complex (referred to as ParBS_F_ hereafter), ParA_F_ exhibits foci formation that occupies the mid-cell, one-quarter or two-thirds positions along the cell length (Niki and Hiraga 1997; Hatano et al. 2007; Hirano et al. 1998).

Recent super-resolution imaging also provides direct evidence of nucleoid localisation of ParA (Le Gall et al. 2016; McLeod et al. 2017). Additional *in vitro* reconstitution on DNA carpets and *in vivo* experiments have led to the idea that chemophoretic gradients generated by dimerisation of ParA upon ATP binding and stimulation of ATPase activity by the partitioning complex drive the directional movement of the replicated plasmids to achieve spatial segregation (Vecchiarelli et al. 2013b, 2014; Hu et al. 2015; Hwang et al. 2013; Vecchiarelli et al. 2010). Furthermore, the elasticity of the bacterial nucleoid itself has been shown to play a role, leading to the proposal of a DNA-relay model (Lim et al. 2014; Surovtsev et al. 2016).

However, *in vitro*, ParA_F_ has also been observed to undergo polymerisation, forming filaments upon interaction with ATP and radial asters in complex with ParBS_F_ ^(^Lim et al. 2005^)^. Such filaments in the presence of ParBS_F_ were also observed *in vivo* in 15 % of the cells. Further, these filaments have been reported to grow at a rate of 0.18 ± 0.05 µm per minute, similar to the rates at which plasmids and chromosomes segregate in bacteria (Lim et al. 2005). Polymeric structures of ParA_F_ in the presence of ATP have also been observed using electron microscopy (Bouet et al. 2007). Moreover, such filaments and ATP-dependent polymers have also been observed in the case of other ParA (Ringgaard et al. 2009; Hui et al. 2010; Volante and Alonso 2015) and related ParA family members like ParF (Barillà et al. 2007, 2005; Schumacher et al. 2012) and Soj (Leonard et al. 2005, 2004). Interestingly, the polymerisation of such filaments in ParA_F_ is mediated only by ATP, wherein non-specific DNA is thought to play a role in inhibiting the formation of such polymers (Bouet et al. 2007). On the contrary, the ParA homolog Soj from *Bacillus subtilis*, *Thermus thermophilus* and ParA2 (a member of the Type-Ia superfamily like ParA_F_) from *Vibrio cholerae* assembles into a higher-order nucleoprotein complex in the presence of DNA (Leonard et al. 2005; Hui et al. 2010; Parker et al. 2021; Leonard et al. 2004; Chodha et al. 2023). Although these early observations supported a cytoskeletal polymerisation-based model for plasmid segregation by ParA_F_, more recent studies from several ParA members, as mentioned above, have suggested mechanisms of DNA partitioning independent of ParA polymerisation. Models like the diffusion ratchet mechanism (Vecchiarelli et al. 2012; Brooks and Hwang 2017; Hatano and Niki 2010; Hu et al. 2015, 2017), DNA relay mechanism (Lim et al. 2014; Surovtsev et al. 2016), and the Hitch-Hiking models (Le Gall et al. 2016) have indeed questioned the physiological relevance of ParA polymerisation.

Interestingly, Pogliano and colleagues had reported a spontaneous mutant of ParA_F_ (M315I Q351H) that stabilised the polymers in *E. coli* cells (Lim et al. 2005). Further, we had earlier carried out an extensive mutational analysis of the C-terminal region in ParA_F_ and found a significant role for the C-terminal helix in plasmid maintenance (Mishra et al. 2022). More recently, a cryo-EM study of filaments formed by ParA from *Vibrio cholerae*, the ParA2_Vc_ protein, has been suggested to involve the C-terminal helix of the protein (Parker et al. 2021). Thus, these recent cryo-EM studies prompted us to revisit the role of polymerisation in ParA function. Here, we report that the C-terminal helix 14 and helix 16 play a significant role in stabilising the SopA polymers. We first show that a single mutation (Q351H) is sufficient to recapitulate the cytoplasmic filaments seen in ParA_F_ M315I Q351H (SopA1). We also report another mutant of ParA_F_, W362E, which assembles into similar polymers in the cytoplasm of *E. coli* cells, suggesting that the mutants act to stabilise the ParA_F_ polymers *in vivo*. We show that while ParA_F_ Q351H and W362E retain their ability to interact with the ParA_F_ promoter region to act as super-repressors, the assembled filaments are not nucleoid-associated. Further, ParA_F_ Q351H strongly induces polymerisation of the wild-type ParA_F_ when co-expressed. Moreover, using various mutants ParA_F_ in the conserved K120 residue in the Walker A motif, we find that polymerisation of ParA_F_ Q351H and W362E requires ATP binding and depends upon the conformational change to the ParA-ATP* state. Finally, we show that polymerisation is completely abolished when we change a charged residue, E375, in the C-terminal helix 16 or R320 in helix 14 to alanine. While ParA_F_ E375A and ParA_F_ R320A retain weak transcriptional activity in response to the ParBS_F_ complex, the mutants fail to maintain plasmids in cells stably. Thus, our studies here highlight a role for helix 14 and the C-terminal helix 16 in polymerisation of ParA_F_ and are suggestive of a requirement of higher order structures (oligomerisation) for co-operative binding of ParA_F_ to DNA and F-plasmid segregation functions.

## RESULTS

### Mutations in ParA_F_ Q351 or W362 stabilise ParA_F_ polymers and impair DNA partitioning

We and others have earlier shown that a C-terminal GFP fusion to ParA_F_ is fully functional for plasmid partitioning, localises to the bacterial nucleoid and forms oscillating foci in the presence of ParBS_F_ complex (Lim et al. 2005; Bouet et al. 2007; Hatano et al. 2007; Ah-Seng et al. 2013; Le Gall et al. 2016; Mishra et al. 2022). ParA_F_-GFP and its variants were expressed from the weakened P_trc_ promoter, while *parBS*_F_ was carried on a mini-F plasmid, pDAG198, that lacked ParA_F_ (*ΔparA_F_, parBS^+^*) under the constitutive P_L*tetO-1*_ promoter (Castaing et al. 2008). In the presence of the *parBS*_F_, the wild type ParA_F_-GFP (SopA) formed foci or appeared as a haze, consistent with earlier reports suggesting nucleoid association **(Fig. 1A)**. On the contrary, ParA_F_ M315I Q351H-GFP (called SopA1) formed filament structures in the cell **(Fig. 1A)** as reported previously (Lim et al. 2005). To test if the Q351H mutation, instead of the double mutations, was sufficient to trigger the polymerisation of ParA_F_, we generated ParA_F_ Q351H-GFP and expressed it in *E. coli* from the same weakened P*_trc_* promoter. ParA_F_ Q351H-GFP also assembled into cytoplasmic filaments similar to that formed by ParA_F_ M315I Q351H-GFP, showing that the mutation in the residue Q351 was sufficient to induce polymerisation of ParA_F_ **(Fig. 1A).** During the course of our studies on the role of the C-terminal helix on ParA_F_ function (Mishra et al. 2022), we identified another mutation, W362E, which likewise resulted in the assembly of ParA_F_-GFP into cytoplasmic filaments **(Fig. 1A)**. In the case of ParA_F_ M315I Q351H, 56.5 % ± 11.27 (95% CI, n=3, at least 100 cells were counted for each experiment) formed cytoplasmic filaments. Further, 41.3 % ± 14.8 (95% CI, n=3; at least 100 cells were counted for each experiment) of the cells in the case of ParA_F_ Q351H contained filaments, while in the case of ParA_F_ W362E, 38.0 % ± 17.9 (95% CI, n=3; at least 100 cells were counted for each experiment) of the cells contained filaments in presence of *parBS*_F_ complex **(Fig. 1B)**.

**Figure 1.**
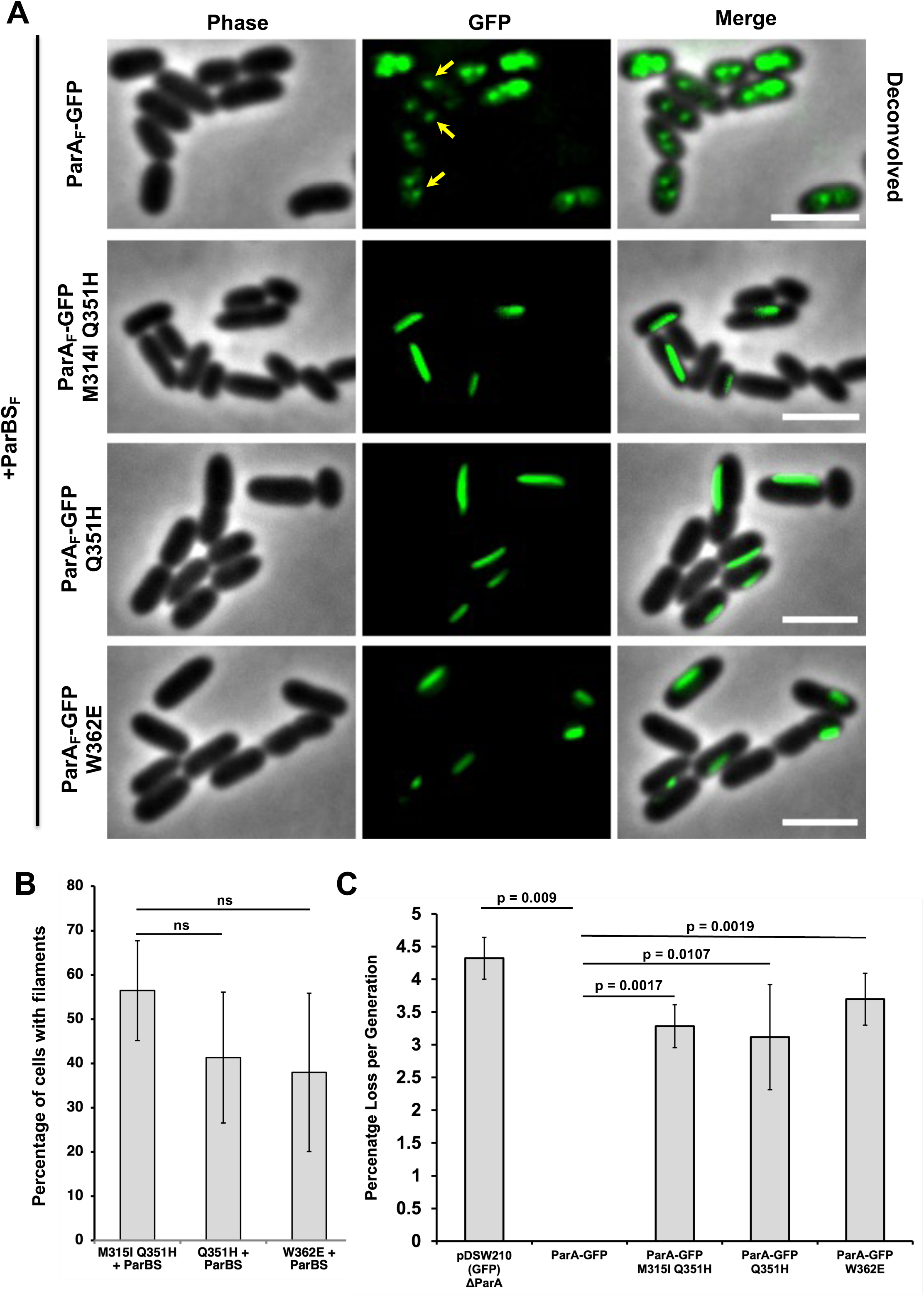
ParA_F_ Q351H and ParA_F_ W362E assemble into cytoplasmic filaments in *E. coli*. **(A)** In the presence of the ParBS complex, wild-type ParA_F_-GFP appears as a haze or forms foci. ParA_F_ M315I Q351H-GFP (SopA1-GFP), ParA_F_ Q351H-GFP and ParA_F_ W362E assemble into linear cytoplasmic filaments. ParA_F_ and the various mutants described were expressed from the weak P*_trc_* promoter in pDSW210 as C-terminal GFP fusions by the addition of 100 µM IPTG for 2 hours in *E. coli* MC4100. Cultures were grown to an OD_600_ of 0.2 before the addition of IPTG for protein expression. A mini-F plasmid carried the *parBS* locus under the constitutive P*_LtetO_* promoter and lacked ParA_F_ (*ΔparA*_F_ *parBS^+^*). **(B)** Quantification of percentage of cells carrying ParA_F_ M315I Q351H-GFP (SopA1-GFP), ParA_F_ Q351H-GFP and ParA_F_ W362E-GFP filaments. Percentage of cells having filaments were 56.5 % ± 11.27, 41.3 % ± 14.8 and 38.0 % ± 17.9 (95% CI, n=3; number of cells counted > 100 for each experiment) in the case of ParA_F_ M315I Q351H-GFP, ParA_F_ Q351H-GFP and ParA_F_ W362E-GFP, respectively. **(C)** Quantification of loss of the mini-F plasmids in cultures carrying wild-type ParA_F_-GFP, ParA_F_ M315I Q351H-GFP (SopA1-GFP), ParA_F_ Q351H-GFP and ParA_F_ W362E-GFP. While ParA_F_-GFP maintained the plasmids stably, the loss of plasmids in all ParA_F_ mutants was similar to that of the vector control (lacking ParA_F_ but expressing GFP alone). The partitioning complex ParBS (where indicated) was provided from a mini-F plasmid (pDAG198; *ΔparA_F_ parBS*^+^) that lacked ParA_F_ but carried the *parBS* locus under the constitutive P*_LtetO_* promoter. The same two plasmids were used for the plasmid stability assays in **(A)**. Cells were mounted onto a 1.6% LB agarose slide and imaged using an epifluorescence microscope. A two-sided Student’s -t-test was used to calculate P-values in case of **(B)**, and since we expect the plasmid loss in ParA_F_ mutants to be only higher than in ParA_F_, a one-sided one-sample t-test was used to assess the significance in this case **(C)**. ns represents a p-value > 0.05. All experiments were repeated at least thrice (n = 3, three biological replicates). Error bars in **(B)** are inferential and represent the 95% confidence interval and the error bars in **(C)** represent the standard deviation (SD) of the mean. Scale bars represent 3 µm for all images.

The cytoplasmic filaments assembled by ParA_F_ M315I Q351H-GFP have been reported to be less dynamic as compared to the WT-ParA_F_ and fail to support F plasmid maintenance (Lim et al. 2005). We, therefore, tested whether the assembly of ParA_F_ Q351H and W362E into polymers also resulted in the loss of mini-F plasmids from cells and carried out plasmid stability assays using the two-plasmid system (Ah-Seng et al. 2013; Mishra et al. 2022^)^. ParA_F_-GFP and its variants were expressed from the weakened P_trc_ promoter and a mini-F plasmid lacking ParA_F_ but containing *parBS_F_* under the constitutive P_LtetO-1_ promoter (mini-F Cam^R^ *P_LtetO-1_::ΔparA_F_*, *parBS*^+^; Castaing ^e^t al. 2008^)^ constituted the second plasmid whose maintenance was tested. As would be expected, in the presence of wild-type ParA_F_-GFP, mini-F plasmids were maintained stably, whereas a loss rate of 4.3 % ± 0.32 (SD, n=3; at least 400 colonies were tested for each experiment) per generation was observed in the absence of ParA_F_. The mini-F plasmid loss rates in cultures carrying ParA_F_ mutants Q351H and W362E were 3.1 % ± 0.80 (SD, n=3; at least 400 colonies were tested for each experiment) and 3.7 % ± 0.40 (SD, n=3; at least 400 colonies were tested for each experiment) respectively, which was comparable to those lacking ParA_F_ **(Fig. 1C)**. ParA_F_ M315I Q351H-GFP (SopA1) also exhibited plasmid loss from the cultures as reported earlier (Lim et al. 2005) at the rate of 3.3 % ± 0.33 (SD, n=3; at least 400 colonies were tested for each experiment), showing that these ParA_F_ mutants that assemble into stable polymers are impaired in partitioning plasmids.

### ParA_F_ Q351H and ParA_F_ W362E or W362A assemble into polymers in the absence of *parBS*_F_

Earlier reports have suggested that ParB influences the polymerisation of ParA_F_ ^(^Bouet et al. 2007^)^. We thus tested if ParA_F_ M315I Q351H, ParA_F_ Q351H and ParA_F_ W362E assembled into cytoplasmic filaments in the absence of *parBS*_F_. Cytoplasmic filaments of ParA_F_ M315I Q351H-GFP, ParA_F_ Q351H-GFP and ParA_F_ W362E-GFP were observed in the absence of ParBS_F_ as well **(Fig. 2A)**, showing that the polymerisation of these ParA_F_ mutants were independent of the partitioning complex. We also substituted W362 with a non-polar amino acid, alanine (W362A) and found that ParA_F_ W362A-GFP also assembled into polymers similar to those formed by ParA_F_ M315I Q351H, ParA_F_ Q351H, and ParA_F_ W362E **(Fig. 2A)**. Moreover, in the absence of *parBS*_F_, quantification of the number of cells with filaments showed no significant differences between the various ParA_F_ mutants **(Fig. 2B)**. While cells expressing ParA_F_ M315I Q351H-GFP showed 39.2 % ± 3.37 (95% CI, n=3; at least 100 cells were counted for each experiment) of cells containing filaments, ParA_F_ Q351H-GFP and ParA_F_ W362E-GFP resulted in 48.7 % ± 9.46 (95% CI, n=3; at least 150 cells were counted for each experiment) and 42.80 % ± 8.23 (95% CI, n=3; at least 150 cells were counted for each experiment) of cells having filaments, respectively **(Fig. 2B)**. The percentage of cells having filaments in the case of ParA_F_ W362A was 53.8 % ± 15.12 (95% CI, n=3; at least 150 cells were counted for each experiment) **(Fig. 2B)**. Further, ParA_F_ Q351H and W362E filaments showed a mean length of 0.88 µm ± 0.19 (95% CI, n=3; number of cells counted were > 73 and < 121) and 0.71 µm ± 0.14 (95% CI, n=3; number of cells counted were > 59 and < 99) respectively (**Fig. 2C**).

**Figure 2.**
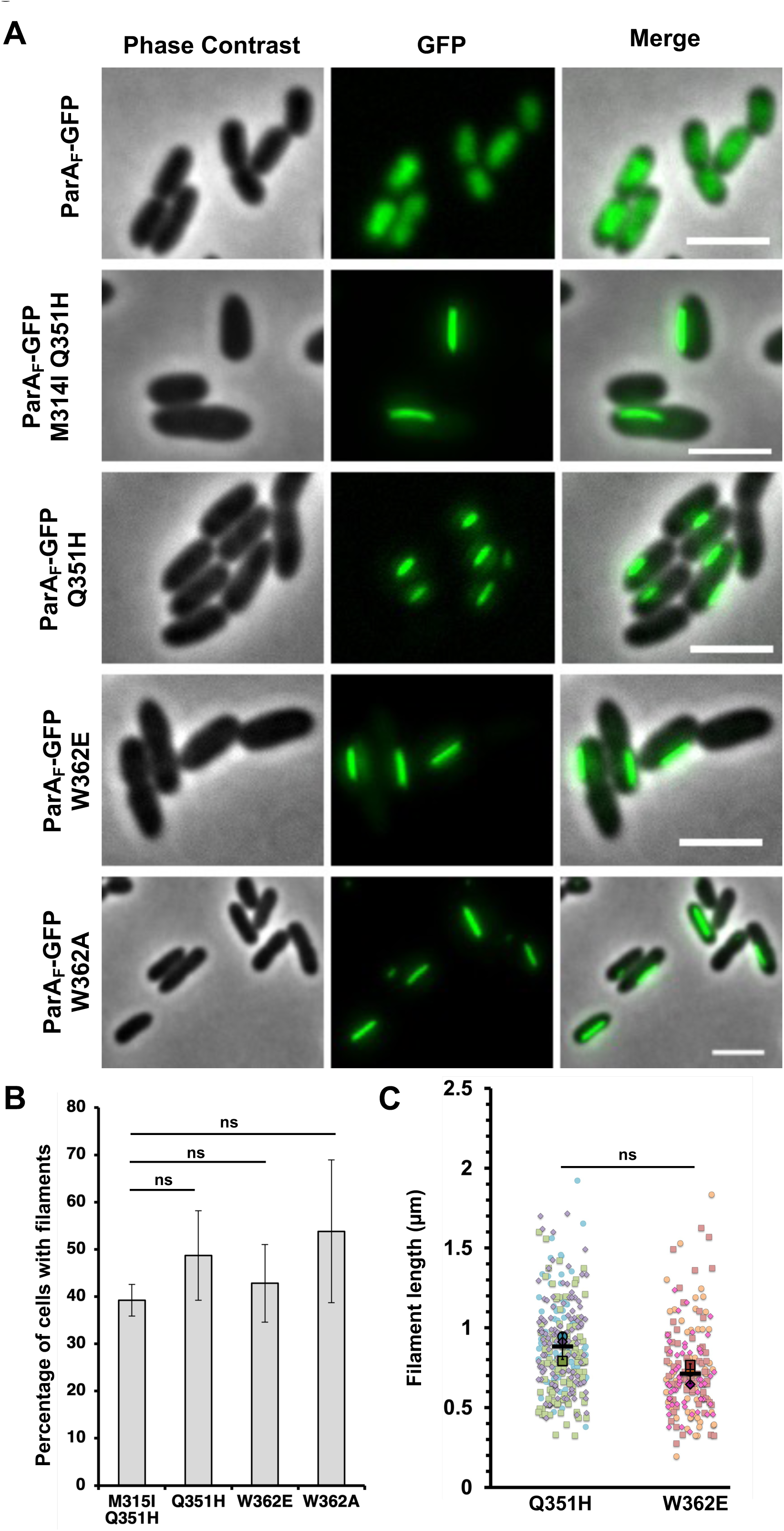
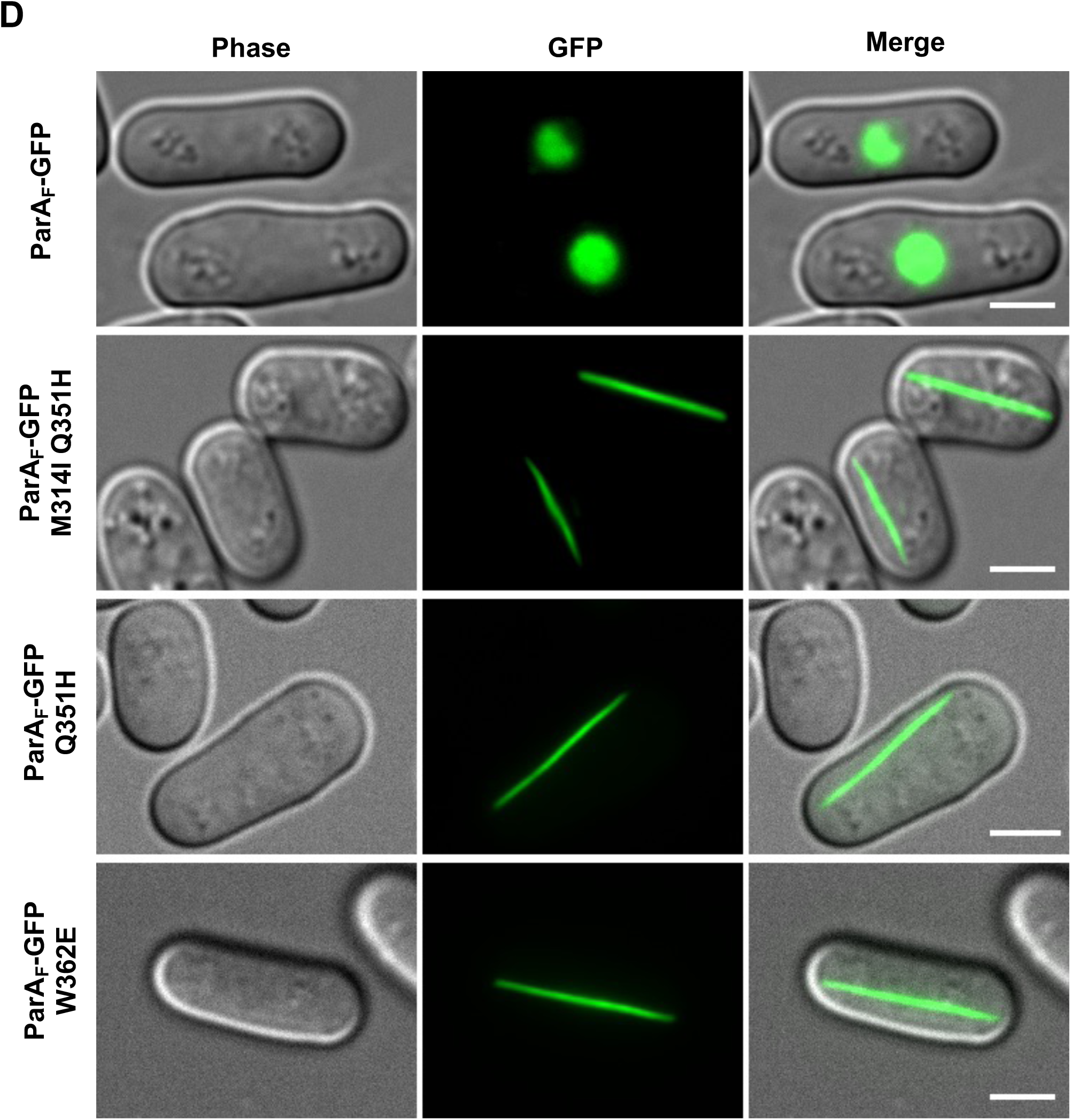
Assembly of ParA_F_ Q351H, ParA_F_ W362E or ParA_F_ W362A into cytoplasmic filaments is independent of the *parBS* partitioning complex. **(A)** In the absence of the *parBS* complex, ParA_F_-GFP does not form foci but appears to be over the nucleoid. ParA_F_ M315I Q351H-GFP (SopA1-GFP), ParA_F_ Q351H-GFP, ParA_F_ W362E-GFP and ParA_F_ W362A-GFP assemble into straight cytoplasmic filaments, even in the absence of the *parBS* complex. **(B)** The percentage of cells having cytoplasmic filaments was similar in the case of ParA_F_ M315I Q351H-GFP [39.2 % ± 3.37 (95% CI, n=3; number of cells counted > 100 for each experiment)], ParA_F_ Q351H-GFP [48.7 ± 9.5 (95% CI, n = 3; number of cells counted > 150 for each experiment)], ParA_F_ W362E-GFP [42.8 ± 8.2 (95% CI, n = 3; number of cells counted > 150 for each experiment)] and ParA_F_ W362A-GFP filaments [53.8 ± 15.1 (95% CI, n = 3; number of cells counted > 150 for each experiment)]. **(C)** Superplots showing the quantification of the length of the cytoplasmic filaments assembled by ParA_F_ Q351H-GFP and ParA_F_ W362E-GFP. No significant difference was observed with an average length of 0.88 µm ± 0.08 (SD, n = 3; number of cells counted > 73 < 121) in the case of ParA_F_ Q351H-GFP and 0.71 µm ± 0.06 (SD, n = 3; number of cells counted > 59 < 99) in the case of ParA_F_ W362E-GFP. ParA_F_ and the various mutants described were expressed from the weak P_trc_ promoter in pDSW210 as C-terminal GFP fusions by the addition of 100 µM IPTG for 2 hours in *E. coli* MC4100. Cultures were grown to an OD_600_ of 0.2 before the addition of IPTG for protein expression. **(D)** *S. pombe* cultures carrying pREP42-ParA_F_-GFP or ParA_F_ mutants M315I Q351H, Q351H or W362E were grown in EMM medium in the absence of thiamine for 20 – 24 hours to allow for the expression of the ParA_F_-GFP protein. ParA_F_ localised to the nucleus (possibly by diffusion and retention due to DNA binding activity), and ParA_F_ M315I Q351H-GFP (SopA1-GFP), ParA_F_ Q351H-GFP and ParA_F_ W362E-GFP assembled into polymers in the cytoplasm. Bacterial cells or yeast cells were mounted onto a 1.6% agarose slide and imaged using an epifluorescence microscope. A two-sided Student’s -t-test was used to calculate P-values in the case of **(B and C)**. ns represents a p-value > 0.05. All experiments were repeated at least thrice (n = 3, three biological replicates). Error bars are inferential and represent the 95% confidence interval in **(B)** and represent the standard deviation (SD) of the mean in **(C)**. Scale bars represent 3 µm for all images.

Fission yeast (*Schizosaccharomyces pombe*) has been used as a heterologous host to study the assembly of cytoskeletal proteins of bacteria and chloroplast (Srinivasan et al. 2007, 2008; Pande et al. 2022; TerBush et al. 2018; TerBush and Osteryoung 2012). We thus tested if these ParA_F_ mutants (M315I Q351H, Q351H and W362E) assembled to filaments in fission yeast as well. We expressed ParA_F_ M315I Q351H, ParA_F_ Q351H or ParA_F_ W362E as C-terminal GFP fusions in fission yeast using the medium-strength thiamine repressible promoter *nmt41/42* (Basi et al. 1993). We observed that, while ParA_F_-GFP was localised to the nucleus, probably owing to the nsDNA binding activity, ParA_F_ M315I Q351H-GFP, ParA_F_ Q351H-GFP and ParA_F_ W362E-GFP assembled into polymeric structures in the cytoplasm of *S. pombe* **(Fig. 2D)**. Interestingly, the filament length spanned the length of *S. pombe* cells, suggesting that the limited filament length observed in *E. coli* was possibly a result of either the physical limit of cell length or the amount of protein available for polymerisation. These observations further suggest that the polymerisation of these mutants was intrinsic to ParA_F_ and that these filaments were formed independent of any host factor specific to assembly in bacteria. Taken together, these results suggest that the mutations in the residues Q351 and W362 result in the formation of stable polymers of ParA_F_, and the assembly into cytoplasmic filaments is independent of the ParBS_F_ partitioning complex.

### Q351 and W362 influence non-specific DNA binding

ParA_F_-mediated plasmid partitioning is majorly dependent on the nucleoid association of ParA_F_-ATP dimers and the formation of the chemophoretic gradient through which the ParB-*parS* plasmid complex migrates. We had earlier shown that the last C-terminal helix played a role in nsDNA binding and plasmid partitioning. Both Q351H and W362E in ParA_F_ map to the C-terminal region, and it was thus imperative for us to test if these filament-forming mutants were also impaired in interaction with non-specific DNA. The localisation of the filaments formed by these mutant proteins was thus determined by live-cell imaging in conjunction with the nucleoid stain DAPI and the membrane marker FM 4-64. ParA_F_ Q351H-GFP and ParA_F_ W362E-GFP filaments often seemed to localise between the nucleoid and the membrane and appeared not to be co-localised with DNA (**Fig. 3A**). Further 3D structured illumination microscopy (3D-SIM) of cells expressing ParA_F_ Q351H-GFP **(Fig. 3B i)** and ParA_F_ W362E-GFP **(Fig. 3B ii)** showed that the spatial localisation of the polymers were clearly distinct from the nucleoid, suggesting that these filaments were cytoplasmic and were not associated with the nucleoid. **(Fig. 3B i and ii)**. In order to better visualise DNA-free regions and determine if the polymers were indeed localised to DNA-free regions and not bound to the nucleoid, we inhibited cell division using cephalexin for a short duration of 30 minutes and condensed nucleoids using chloramphenicol, stained them with DAPI and carried out live-cell imaging. The ParA_F_ Q351H-GFP and ParA_F_ W362E-GFP filaments were invariably localised to the nucleoid-free regions of the cell **(Fig. 3C)**, and we did not find any co-localisation of filaments with the nucleoid. On the contrary, ParA_F_-GFP was seen to be entirely co-localised with the nucleoid, as expected **(Fig. 3C)**.

**Figure 3.**
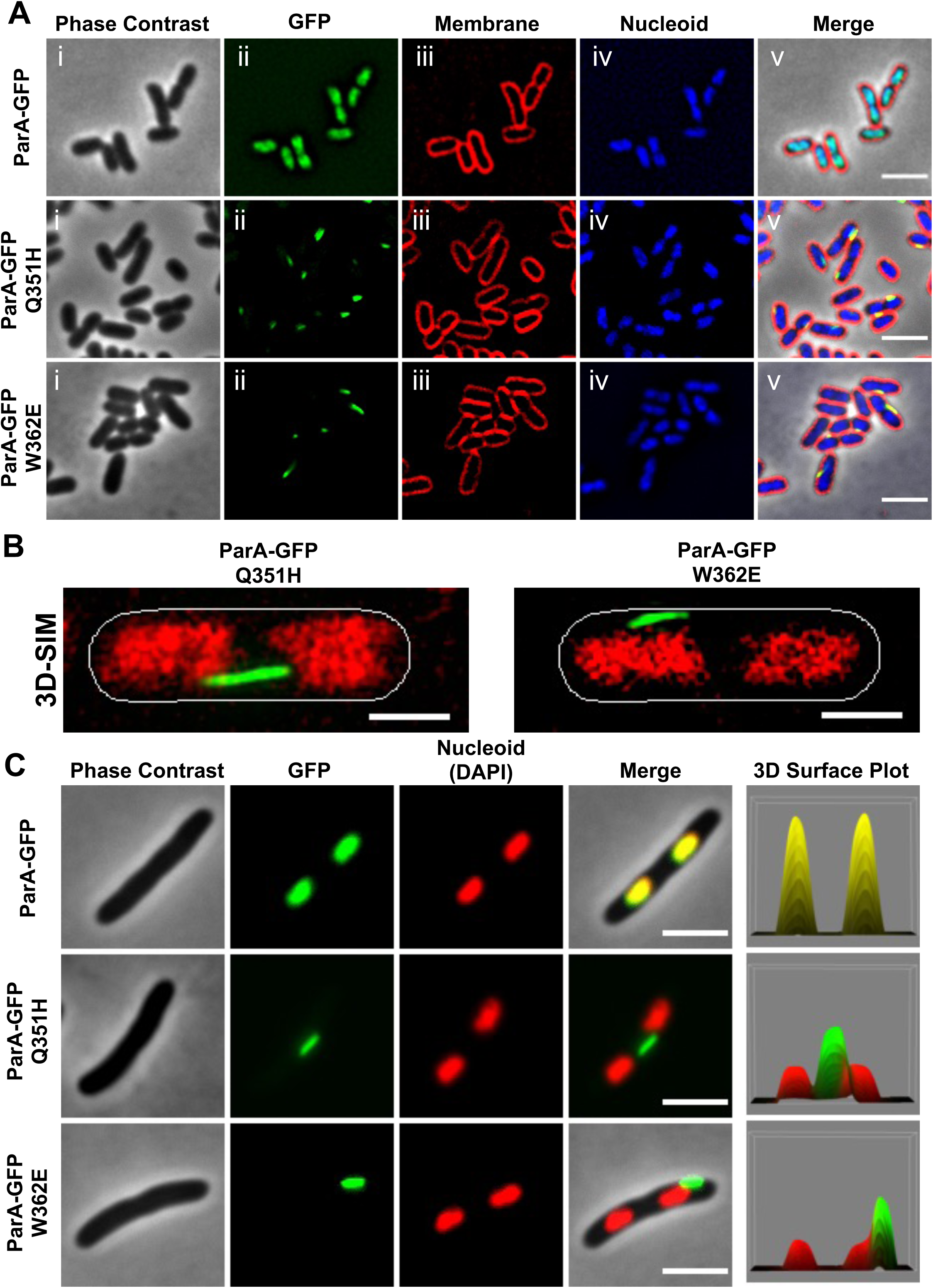
Residues Q351 and W362 influence non-specific DNA binding by ParA_F_. **(A)** Filaments formed by ParA_F_ Q351H-GFP and ParA_F_ W362E-GFP are not nucleoid-associated. The ParA_F_ Q351H-GFP and ParA_F_ W362E-GFP filaments appeared to be localised in the space between the nucleoid and the inner membrane, while ParA_F_-GFP localised over the nucleoids. Nucleoids were stained with DAPI, and the plasma membrane was stained using FM4-64. **(B)** Super-resolution imaging using 3D structured illumination microscopy (3D-SIM) showing the filaments assembled by **(i)** ParA_F_ Q351H-GFP and **(ii)** ParA_F_ W362E-GFP are not associated with the nucleoid. DNA was stained with DAPI and pseudo-coloured red. Cell outlines were obtained from the phase contrast images. **(C)** Filaments formed by ParA_F_ Q351H-GFP and ParA_F_ W362E-GFP are found in the nucleoid-free spaces in the cytoplasm of *E. coli*. Wild-type ParA_F_-GFP localised to the nucleoid. Nucleoids were visualised by staining with DAPI and pseudo-coloured red in the images. Cells were treated with 25 µg/ mL cephalexin for 30 minutes to prevent cell division and 100 µg/ mL chloramphenicol for 10 minutes to condense the nucleoids so that the nucleoid-free space appears pronounced. Scale bars represent 3 µm in all images except in **(B)**, where the scale bar is 1 µm.

### Co-expression of filament-forming mutants can induce polymerisation of wild-type ParA_F_-GFP

Earlier studies have shown that ParA_F_ M315 Q351H (SopA1) can co-assemble with ParA_F_-GFP and induce the formation of ParA_F_-GFP filaments. To further test if ParA_F_ Q351H and ParA_F_ W362E can similarly induce polymerisation of wild-type ParA_F_-GFP, which are otherwise localised to the nucleoid, we resorted to in vivo imaging of a two-plasmid system. We expressed the wild-type ParA_F_-GFP fusion from the *P_trc_* promoter (pDSW210 vector, Amp^R^) as described above and simultaneously expressed versions of ParA_F_ Q351H or ParA_F_ W362E lacking the fluorescent protein tag from the highly regulated arabinose promoter (pBAD33 vector, Cm^R^). Upon co-expression of untagged wild-type ParA_F_ from the pBAD33 vector, ParA_F_-GFP was found localised to the nucleoid as expected **(Fig. 4A)**. However, ParA_F_-GFP assembled into polymers in cells co-expressing untagged ParA_F_ Q351H **(Fig. 4A)**, suggesting that ParA_F_ Q351H co-assembled with wild-type ParA_F_-GFP and stabilised ParA_F_ in its polymeric form *in vivo*. On the contrary, ParA_F_ W362E only weakly induced polymerisation of ParA_F_-GFP and most of the ParA_F_ remained nucleoid-associated **(Fig. 4A)**. Further, the polymers assembled by wild-type ParA_F_-GFP were more clearly visualised upon preventing cell division by treatment of cells with cephalexin (**Fig. 4B)**. Thus, using co-expression of WT-ParA_F_-GFP and untagged versions of ParA_F_ mutants and imaging, we show that the mutants ParA_F_ Q351H and ParA_F_ W362E can co-polymerise with wild-type ParA_F_ and sequestered into the polymeric form. Our results here further support the premise that wild-type ParA_F_ has the property to assemble into higher-order oligomers, and these mutations (Q351H and W362E) resulted in the polymerisation of ParA_F_ *in vivo*.

**Figure 4.**
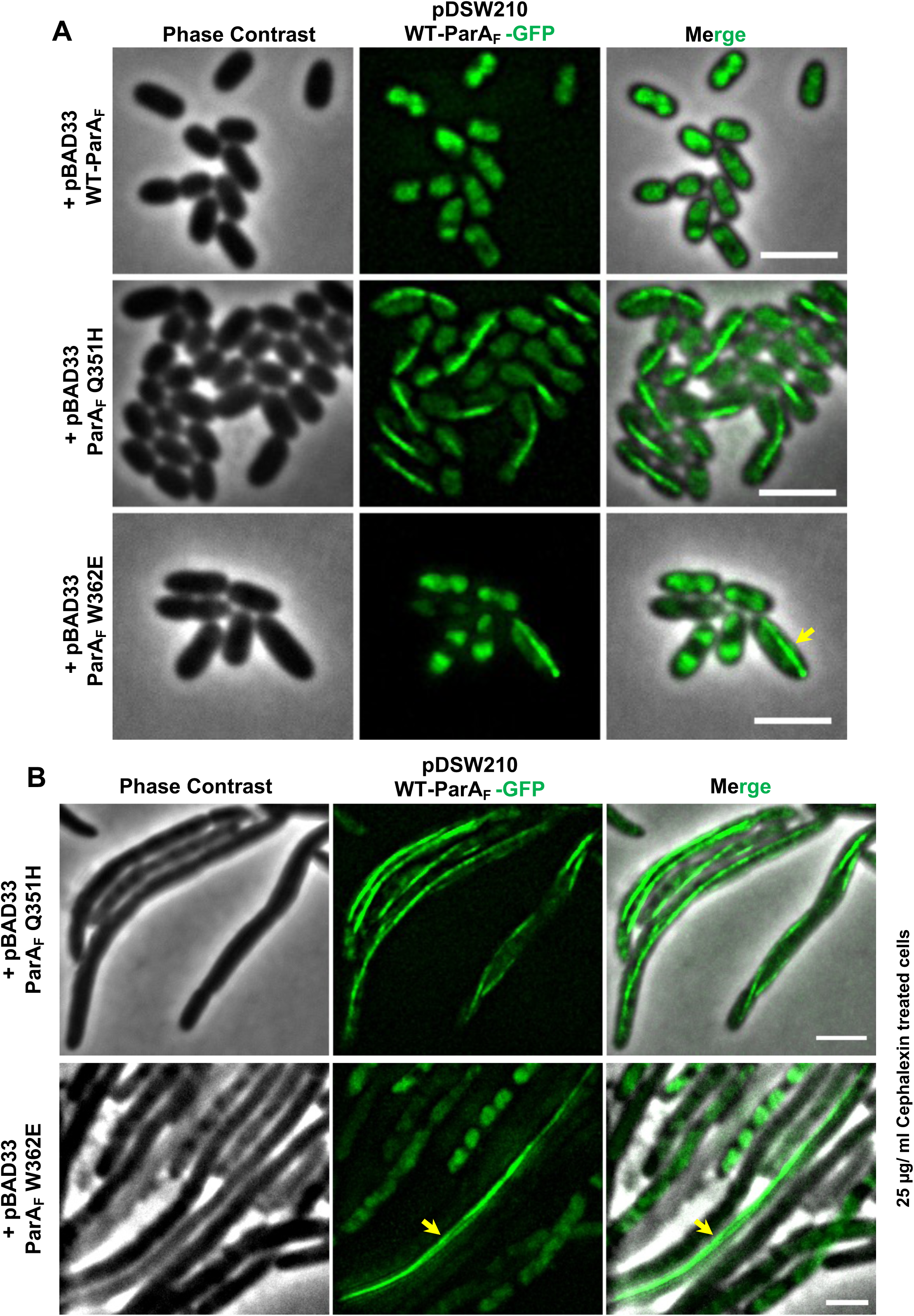
Filament-forming mutants of ParA_F_ can induce the assembly of wild-type ParA_F_-GFP into cytoplasmic filaments. **(A)** Polymerisation of ParA_F_-GFP upon co-expression of untagged ParA_F_ Q351H or ParA_F_ W362E. ParA_F_-GFP localises to the nucleoid upon co-expression with untagged ParA_F_, but co-expression of untagged ParA_F_ Q351H induces the assembly of ParA_F_-GFP into cytoplasmic filaments. Untagged ParA_F_ W362E can also induce the polymerisation of ParA_F_-GFP but is very inefficient compared to ParA_F_ Q351H, and ParA_F_-GFP remains predominantly nucleoid localised. **(B)** Assembly of ParA_F_-GFP into long filaments in cephalexin-treated cells upon co-expression of untagged ParA_F_ Q351H. ParA_F_-GFP was expressed from the weak P_trc_ promoter of pDSW210 vector by the addition of 100 µM IPTG, and untagged versions of ParA_F_, ParA_F_ Q351H or ParA_F_ W362E were co-expressed from the tightly regulated arabinose inducible promoter P_BAD_ from pBAD33 vector. Cells were treated with 25 µg/ mL cephalexin in **(B)** to obtain filamentous cells and inhibit cell division for better visualisation of ParA_F_ filaments. Scale bars represent 2 µm.

### Continuous protein synthesis is required to maintain ParA_F_ filaments

We next attempted to visualise the polymerisation dynamics of ParA_F_ Q351H and ParA_F_ W362E. However, we could rarely capture the phenomenon of foci/ spot to filament formation in our time-lapse images. The increment in the length could not be discerned, probably owing to limiting protein levels due to the weak promoter used (pDSW210; Weiss et al., 1999) and cell division-induced dilution effects. Therefore, we resorted to cephalexin treatment of cells, which inhibited cell division, to monitor filament growth in the presence of IPTG. We were able to monitor filament elongation in these non-dividing cells and were able to capture the growth of the polymers assembled by ParA_F_ Q351H-GFP **(Fig. 5A)** and ParA_F_ W362E-GFP **(Fig. 5B)** in our time-lapse experiments.

**Figure 5.**
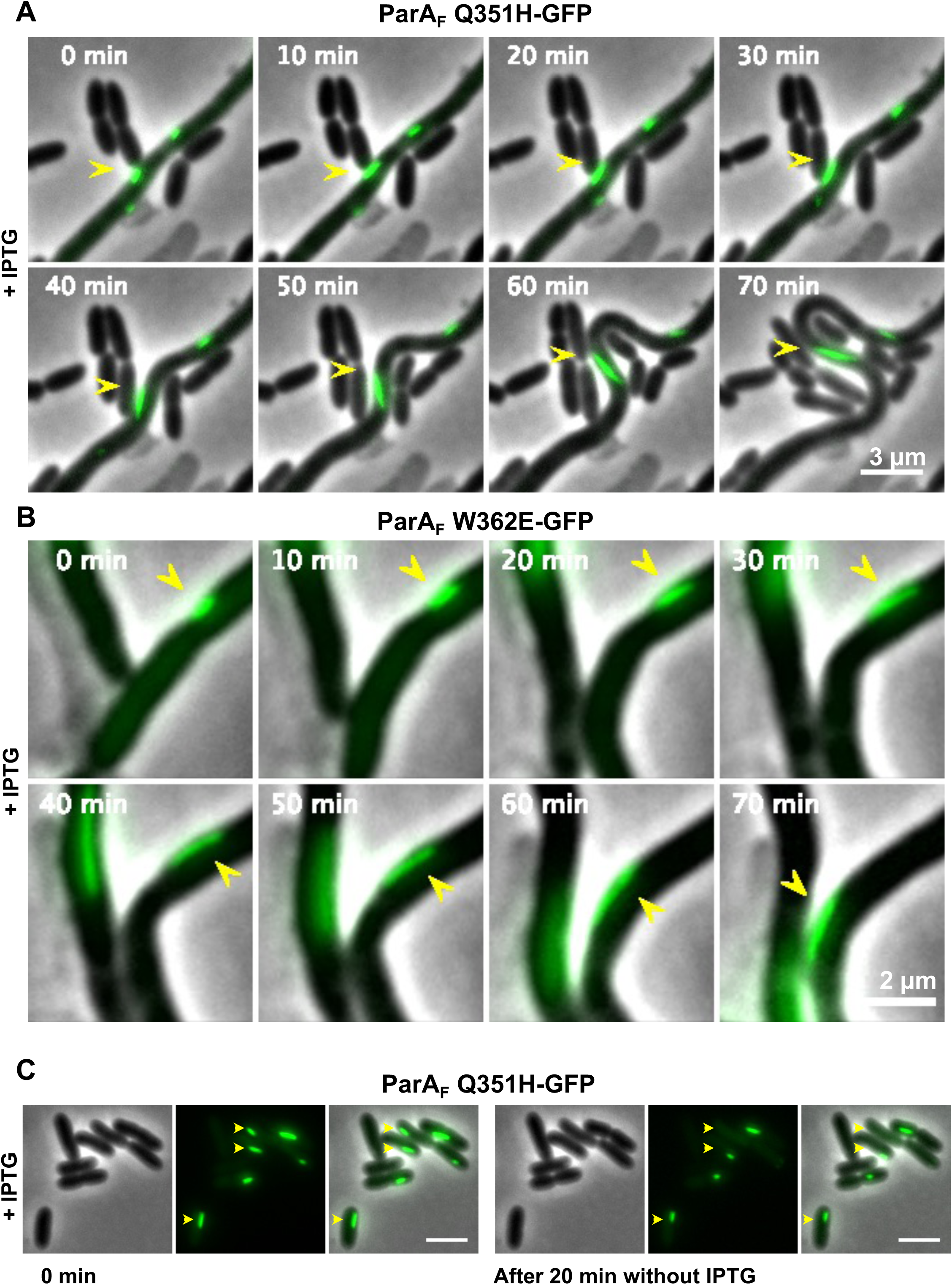
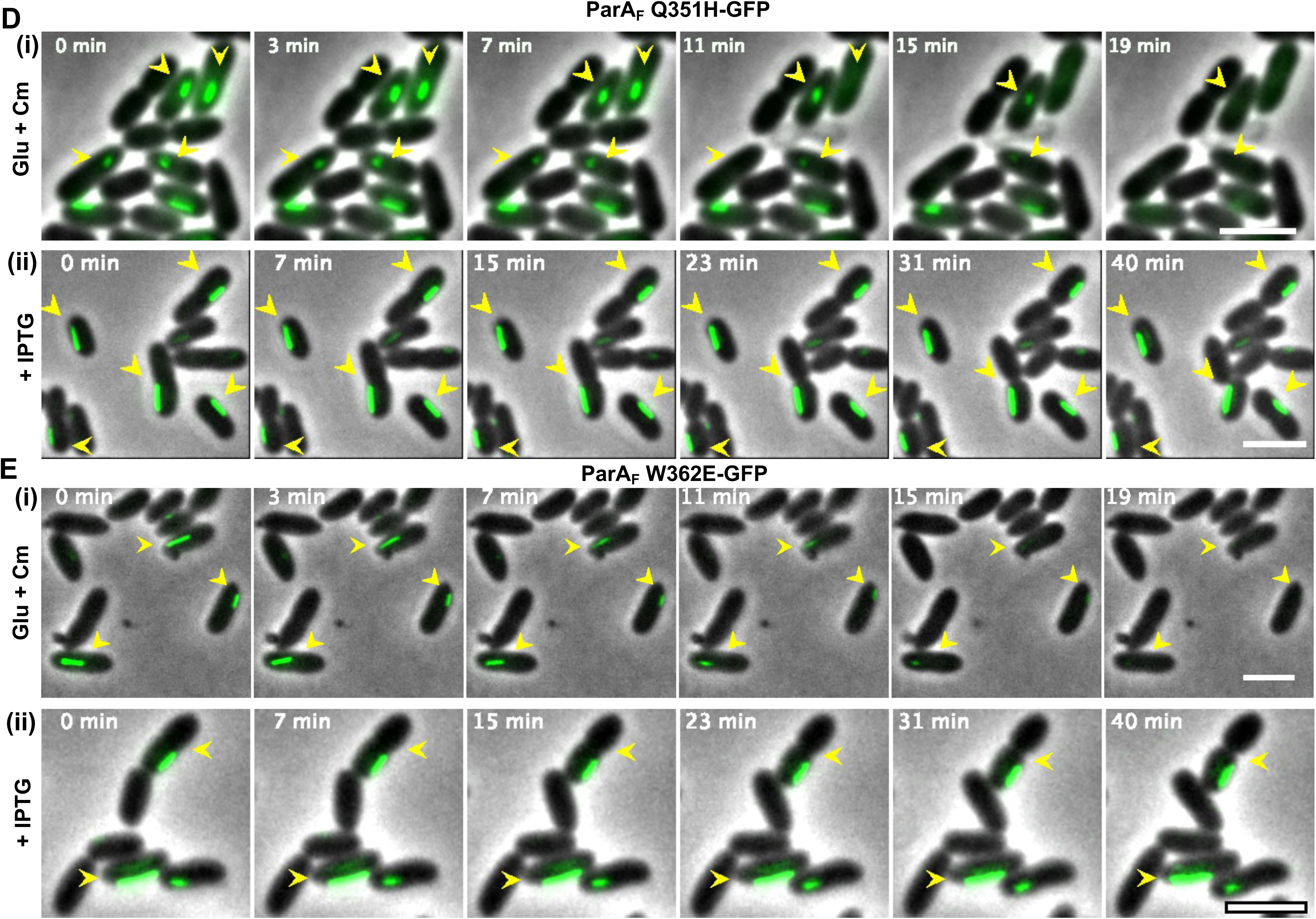
Maintenance of ParA_F_ Q351H and ParA_F_ W362E polymers requires continuous protein synthesis in cells. **(A – B)** Time-lapse images showing the growth of ParA_F_ filaments formed by **(A)** ParA_F_ Q351H-GFP and **(B)** ParA_F_ W362E-GFP. Live cells were imaged on LB-agarose pads containing 100 µM IPTG and 25 µg/ mL cephalexin. Images were acquired every 10 minutes for a total of 70 minutes on a Delta Vision Elite^TM^ microscope. Arrowheads depict the growing filaments. Cells were treated with cephalexin (25 µg/ mL) to prevent cell division to clearly visualise the growth of ParA_F_ filaments. **(C)** ParA_F_ Q351H-GFP filaments appear to shorten in the absence of the inducer IPTG. *E. coli* MC4100 cells carrying pDSW210-ParA_F_ Q351H-GFP were induced 100 µM IPTG and then placed on LB-agarose slides without IPTG. Images were acquired immediately after placing the cells on agarose slides, and the same field was imaged again after 20 minutes. **(D and E)** Time-lapse images showing the depolymerisation of ParA_F_ filaments assembled by **(D i)** ParA_F_ Q351H-GFP and **(E i)** ParA_F_ W362E-GFP in the absence of continued protein synthesis within a period of 20 minutes. Cultures were induced 100 µM IPTG for 2 hours prior to placing on LB-agarose pads and imaging. Chloramphenicol (50 µg/ mL) and 2 % glucose were added to the agarose pads to stop protein synthesis and repress the IPTG inducible promoter in the pDSW210 vector, respectively. Images were acquired every minute for a total of 20 minutes. **(D ii)** Time-lapse images of ParA_F_ Q351H-GFP and **(E ii)** ParA_F_ W362E-GFP showing the stable maintenance of filaments up to 40 minutes in the presence of continuous protein synthesis. Cultures were induced 100 µM IPTG for 2 hours prior to placing on LB-agarose pads and imaging. Cells were imaged on LB-agarose pads containing 200 µM IPTG. Images were acquired every minute for a total of 40 minutes. Scale bars represent 3 µm.

Moreover, we had serendipitously observed that cells containing ParA_F_ W362E-GFP filaments exhibited diffuse cytoplasmic fluorescence when placed on LB agarose pads lacking IPTG for extended periods **(Supplementary Fig. S1**). Further, we also observed that ParA_F_ Q351H-GFP exhibited shortening or loss of ParA_F_ filaments within 20-minutes on LB agarose pads lacking IPTG **(Fig. 5C)**. The depolymerisation of ParA_F_ filaments in the absence of IPTG suggested that the maintenance of polymers required continuous protein synthesis. To determine if the persistence of ParA_F_ filaments depended upon continual protein synthesis, we prevented further induction of protein and blocked protein synthesis in cells expressing ParA_F_ filaments and carried out time-lapse microscopy. We first grew cultures carrying pDSW210-ParA_F_ Q351H or pDSW210-ParA_F_ W362E with IPTG to induce polymerisation. We then carried out time-lapse imaging of these cells placed on IPTG containing agarose pads and compared them with cells imaged on agarose pads containing chloramphenicol and glucose, but lacking IPTG. We added chloramphenicol (Cm), a protein synthesis inhibitor, to prevent further protein synthesis and glucose (Glu) to completely repress ParA_F_ expression from the weakened P*_trc_* promoter. Time-lapse imaging of ParA_F_ Q351H-GFP **(Fig. 5D)** and ParA_F_ W362E-GFP **(Fig. 5E)** in the presence of Glu + Cm **(Fig. 5D i and 5E i)** or IPTG **(Fig. 5D ii and 5E ii)** confirmed that the filaments indeed underwent depolymerisation in the absence of continued protein synthesis within a period of approximately 20 minutes **(Fig. 5 D i and E i)**. These results suggest that both ParA_F_ Q351H and W362E require a critical concentration of protein to assemble into polymers and there exists a cytoplasmic pool of unpolymerised subunits as well in addition to the filaments observed.

### ParA_F_ Q351H and ParA_F_ W362E act as super-repressors of the *parAB_F_* promoter

ParA_F_ is known to weakly repress transcription from its own promoter by binding to the four operator sequences. A winged helix-turn-winged helix (WTH) domain that is distinct from the nsDNA binding residues is responsible for promoter sequence recognition and binding (Hirano et al. 1998; Lemonnier et al. 2000; Libante et al. 2001; Komai et al. 2011; Yates et al. 1999; Mori et al. 1989; Bouet et al. 2007; Ah-Seng et al. 2009; Ravin et al. 2003; Hayes et al. 1994; Abeles et al. 1985). This auto-regulatory activity of ParA_F_ is very weak but is enhanced in the presence of the ParBS_F_ complex. ^(^Ravin et al. 2003; Mori et al. 1989; Hirano et al. 1998; Libante et al. 2001^)^. While the filaments formed by ParA_F_ Q351H and ParA_F_ W362E were not nucleoid-associated, we tested if ParA_F_ Q351H and ParA_F_ W362E (presumably the cytosolic subunits) could at least bind the specific DNA sequences in the promoter region of ParA_F_ and act to repress gene expression. In order to do so, we resorted to using an *in vivo* reporter assay based on *lacZ* fusion to the P*par_F_* promoter and utilised the *E. coli* DLT1127 strain (PparF::lacZ; Ravin and Lane 1999). As reported earlier, we made use of ParA_F_ constructs that carried a stop codon between ParA_F_ and GFP (Mishra et al. 2022) **(Fig. 6A)**. We transformed ParA_F_ or the mutant plasmids into the DLT1127 strain of *E. coli* carrying a mini-F plasmid containing ParBS_F_ region but lacking ParA_F_ (mini-F *ΔparA_F_ parBS*^+^) and assayed for promoter repression by spotting serial dilutions of the cultures on X-Gal + IPTG plates. In the presence of ParBS_F_, although ParA_F_ and both the mutants (Q351H and W362E) repressed the expression of LacZ from the P*par_F_* promoter, the extent of repression by ParA_F_ Q351H and ParA_F_ W362E appeared to be stronger **(Fig. 6B; top panel)**. However, in the absence of the ParBS_F_ complex, while ParA_F_ showed a reduced ability to repress LacZ expression as expected, ParA_F_ Q351H and ParA_F_ W362E continued to strongly repress the expression of LacZ (**Fig. 6B, bottom panel**). Further, quantification of the LacZ expression levels by β-galactosidase assays using ONPG as a substrate (**Figure 9** below) also showed that the repressor function of ParA_F_ Q351H and ParA_F_ W362E were not responsive to the presence of the ParBS_F_ complex. Expression of LacZ was repressed by 74.27 % ± 3.22 (SD, n = 3) and 70.85 % ± 4.95 (SD, n = 3) by ParA_F_ Q351H in the presence and absence of ParBS_F_ complex, respectively. Repression of LacZ by ParA_F_ W362E was weaker and showed 44.60 % ± 8.87 (SD, n = 3) and 44.03 % ± 1.16 (SD, n = 3) in the presence and absence of ParBS_F_ complex, respectively. On the contrary, ParA_F_ showed a significant repression of 74.04 % ± 2.05 (SD, n = 3) in the presence of the ParBS_F_ complex compared to a mere 6.40 % ± 9.83 (SD, n = 3) in its absence. These results suggested that the repression of the ParA_F_ promoter activity by ParA_F_ Q351H and ParA_F_ W362E resulted in a so-called super-repressor activity of ParA_F_ independent of the ParBS_F_ complex.

**Figure 6.**
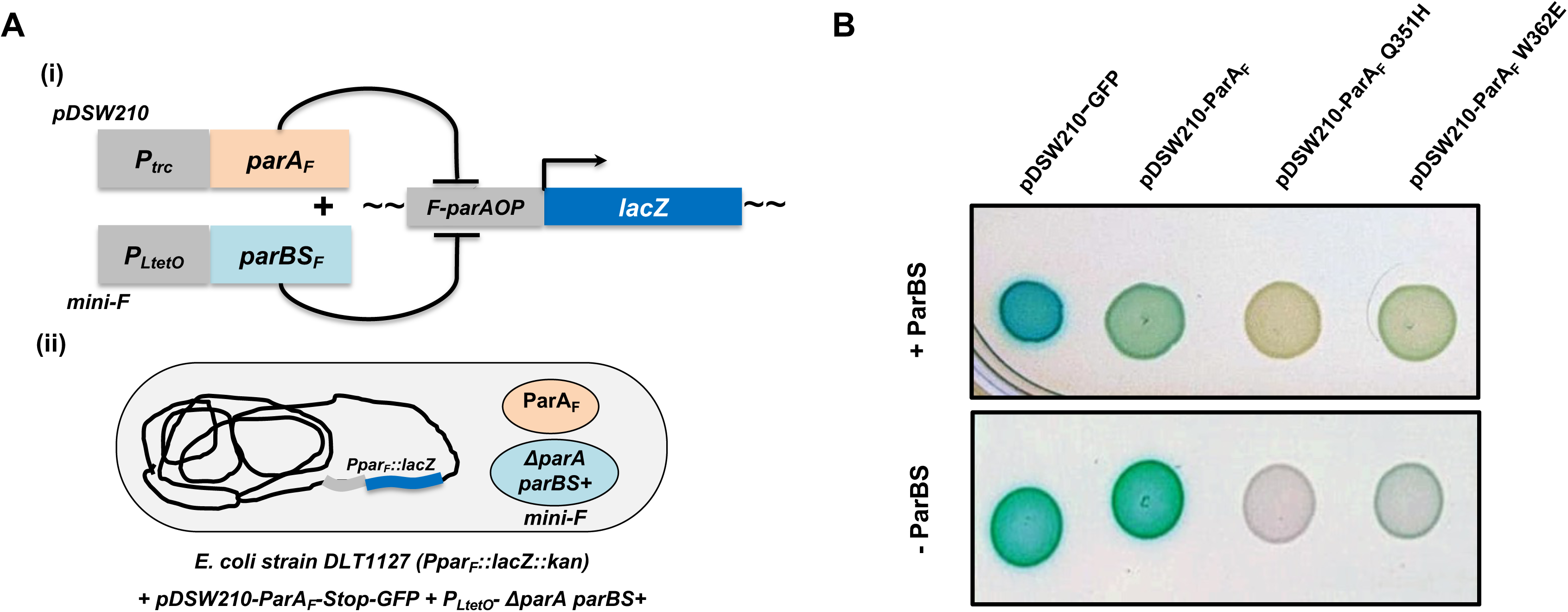
ParA_F_ Q351H and ParA_F_ W362E act as transcriptional repressors of the ParA promoter and are independent of ParB. **(A)** Schematic showing the two-plasmid system and the LacZ-reporter strain used for determining the transcriptional repressor activity of ParA_F_ in the presence or absence of ParBS complex. The strain DLT1127 carried the β-galactosidase gene *lacZ* under the promoter sequences of ParA_F_. ParA_F_ was expressed from the IPTG inducible weak P_trc_ promoter in the pDSW210 vector, and the ParB was expressed from the constitutive promoter P_LtetO_ and was carried on the mini-F plasmid lacking *parA_F_* (*ΔparA parBS^+^*). ParA_F_ acts to strongly repress the expression of *lacZ* from the P_parA_ promoter in the presence of ParB. **(B)** Qualitative β-galactosidase assay showing ParA_F_ Q351H and ParA_F_ W362E acting as super-repressors of the P_parA_ promoter by strongly preventing LacZ expression even in the absence of ParB. While cultures carrying vector control (expressing GFP) did not repress LacZ expression either in the presence or absence of ParBS, ParA_F_ strongly reduced the expression of LacZ only in the presence of the ParBS complex (top panel). ParA_F_ Q351H and ParA_F_ W362E acted independently of ParB and strongly repressed LacZ expression in the absence of ParB as well (bottom panel). Cultures of *E. coli* strain DLT1127 carrying pDSW210-GFP, pDSW210-ParA_F_, pDSW210-ParA_F_ Q351H or pDSW210-ParA_F_ W362E were spotted on LB-agar plates containing appropriate antibiotics, 200 µM IPTG and 40 µM X-Gal and incubated overnight and imaged with CCD colour camera on a phone. Images were colour-corrected using the white-balance plugin in ImageJ v-1.54f.

### Polymerisation is an ATP-dependent process and requires the conformational change to ParA_F_-ATP* state

*In vitro* studies of ParA_F_ and several ParA homologs have shown that polymerisation was strictly an ATP-dependent process (Lim et al. 2005; Leonard et al. 2005; Bouet et al. 2007; Parker et al. 2021). Thus, we wanted to test if the observed *in vivo* polymerisation of ParA_F_ W362E-GFP was dependent upon ATP binding. The residue K120 in ParA_F_ has been reported to be essential for ATP binding and hydrolysis ^(^Libante et al. 2001; Hatano et al. 2007^)^. Changing this residue to glutamic acid (E) in ParA_P1_ has been shown to specifically affect ATP binding (Vecchiarelli et al. 2013a; Chodha et al. 2023; Fung et al. 2001; Davis et al. 1996). We, therefore, generated ParA_F_ double mutants (ParA_F_ K120E W362E-GFP) and assessed their ability to form polymers in *E. coli*. As compared to the single mutant ParA_F_ W362E **(Fig. 7A)**, the double mutation resulted in a complete loss of polymerisation and exhibited diffuse fluorescence of ParA_F_-GFP over the entire cytoplasm of the cell, underlining the role of ATP binding in the polymerisation of these ParA_F_ mutants **(Fig. 7A)**. These results therefore suggest that the assembly of ParA_F_ W362E-GFP into cytoplasmic filaments was strictly an ATP dependent process.

**Figure 7.**
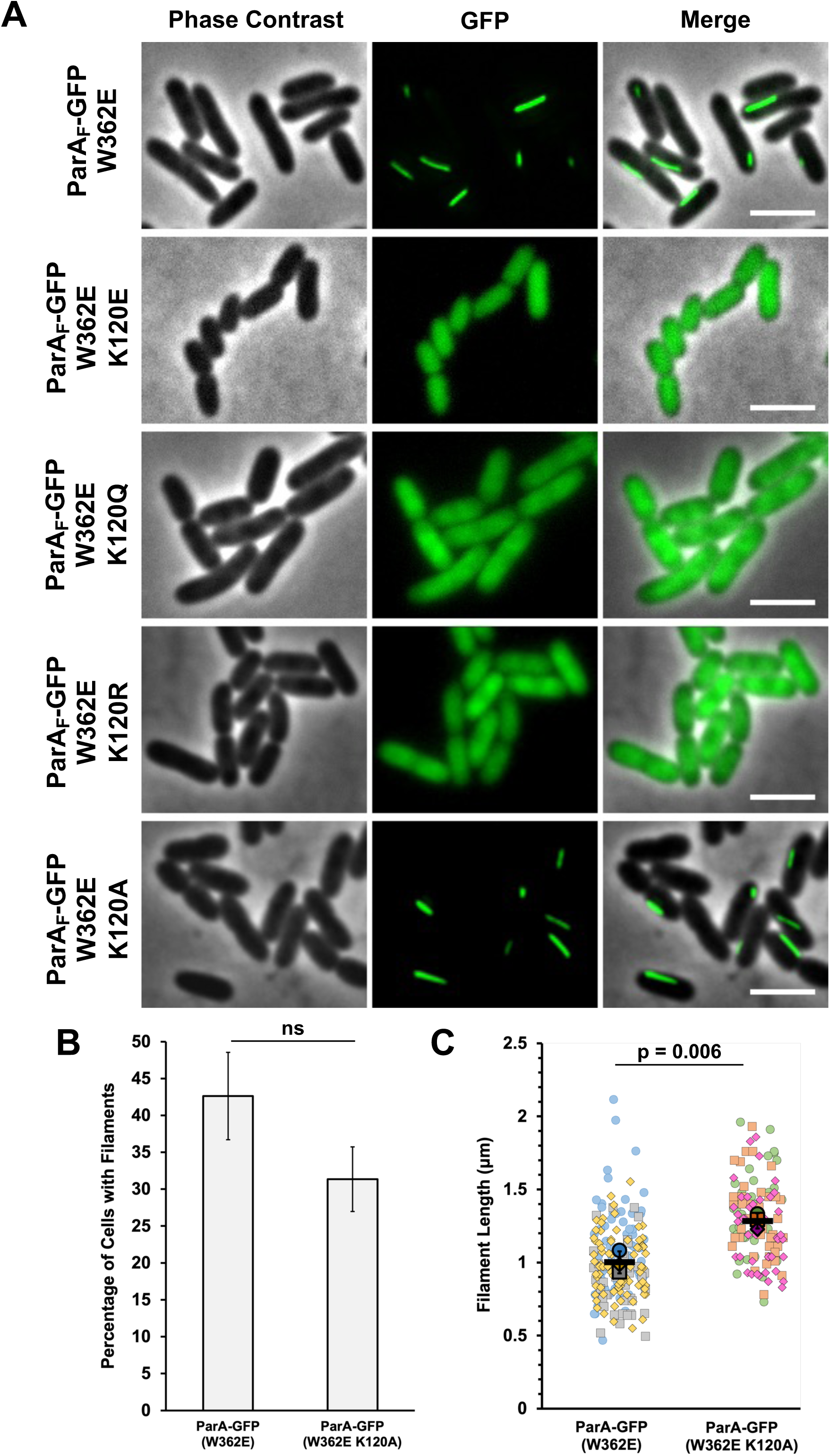
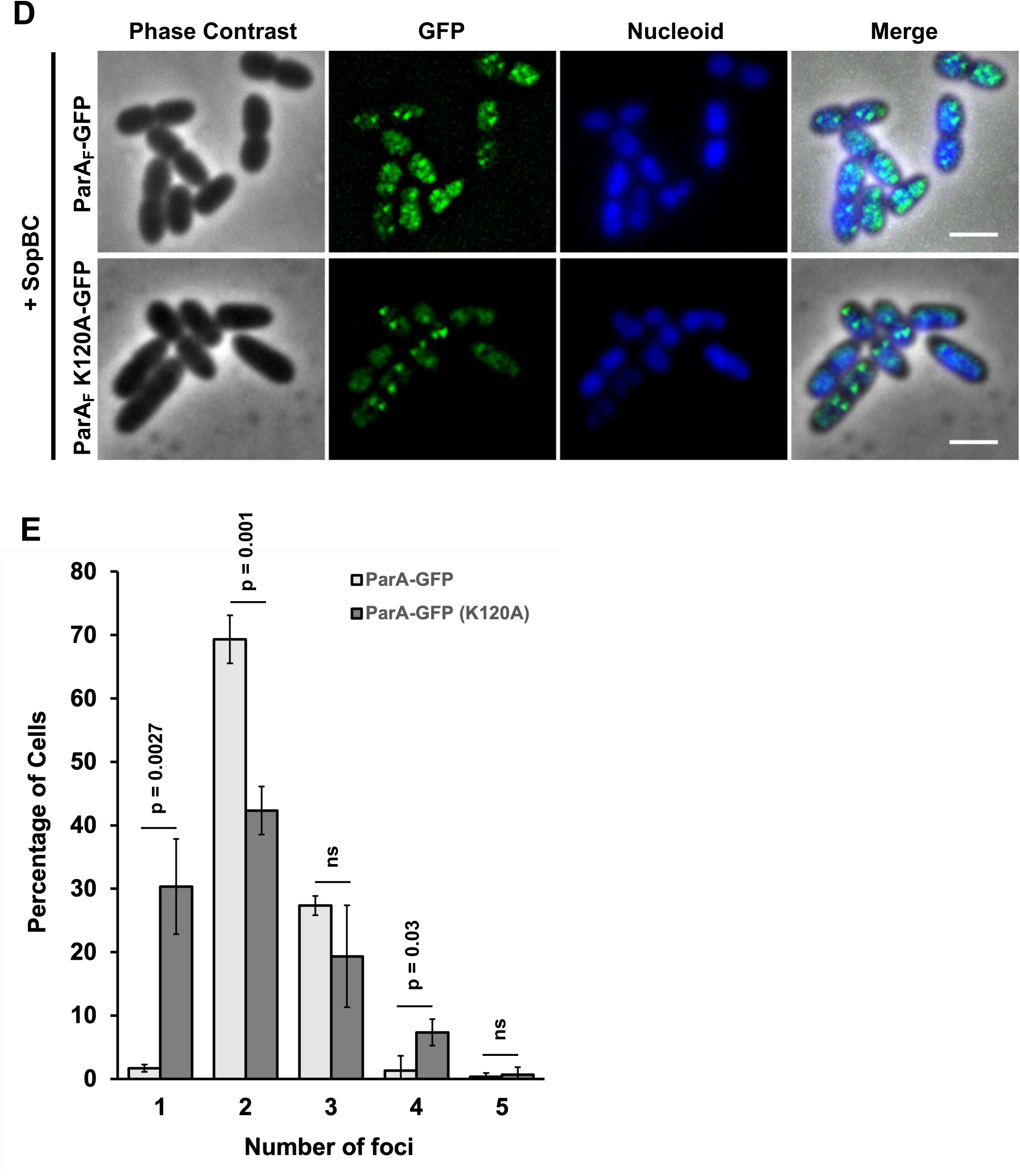
Filament assembly by ParA_F_ W362E-GFP requires ATP-binding and the conformation change to the ParA_F_-ATP* state. **(A)** Images showing the effect of various mutations in the conserved K120 residue on the polymerisation by ParA_F_ W362E-GFP. While ParA_F_ W362E-GFP assembles into cytoplasmic filaments, the introduction of K120E, which abolishes ATP binding in ParA_F,_ fails to form polymers. The introduction of K120R, which retains the ATP binding but does not hydrolyse ATP, also fails to form filaments. However, the introduction of K120Q, which retains the basal ATP hydrolysis, also fails to form filaments. Although ParA_F_ K120R and ParA_F_ K120Q bind the nucleotide, both are defective in achieving the conformational change to ParA_F_-ATP* state upon interaction with ATP, thus showing that ParA_F_-ATP* state is important for the polymerisation of ParA_F_ W362E-GFP. Changing K120 to alanine in ParA_F_ W362E did not affect the filament assembly. **(B)** Quantification of the percentage of cells having cytoplasmic filaments in the ParA_F_-W362E-GFP and ParA_F_-W362E K120A-GFP. While in the case of ParA_F_-W362E-GFP, 42.6 % ± 5.9 (SD, n = 3, at least a total of 150 cells were counted for each experiment) cells had polymers, 31.3 % ± 4.4 (SD, n = 3, at least a total of 150 cells were counted for each experiment) of cells had filaments in the case of ParA_F_-W362E K120A-GFP and were not significantly different. The differences were not statistically significant and had a p-value > 0.05. **(C)** Superplots showing the quantification of the filament lengths in the ParA_F_-W362E-GFP and ParA_F_-W362E K120A-GFP. While in the case of ParA_F_-W362E-GFP, the average length of the filament was 1.0 µm ± 0.08 (SD, n = 3, total number of cells counted were > 75 < 84 for each replicate), ParA_F_-W362E K120A-GFP exhibited an average polymer length of 1.28 % ± 0.05 (SD, n = 3, total number of cells counted were > 40 < 42 for each replicate). The effect size was 0.28 [0.14, 0.43]. **(D)** ParA_F_ K120A-GFP interacts with the partitioning complex, ParBS and forms cloud-like structures and foci over the bacterial nucleoid similar to ParA_F_-GFP, suggesting that K120A mutation allows the conformational change to the ParA_F_-ATP* state and does not affect ATP-binding or nsDNA binding of ParA_F_. **(E)** Quantification of the number of foci assembled per cell by ParA_F_-GFP and ParA_F_ K120A-GFP. A greater percentage of cells have single-foci in the case of ParA_F_ K120A-GFP [30.3 ± 7.5 (SD, n = 3, a total of 100 cells were counted for each replicate)] as compared to ParA_F_-GFP [1.7 ± 0.6 (SD, n = 3, a total of 100 cells were counted for each replicate)]. ns represents non-significant. The error bars in the graphs represent the standard deviation of the mean (SD), and the scale bars in the images represent 2 µm.

ParA_F_ has been proposed to undergo a conformational state change upon ATP-binding (Libante et al. 2001; Vecchiarelli et al. 2010; Zhang and Schumacher 2017; Caccamo et al. 2020; Dobruk-Serkowska et al. 2012; Baxter et al. 2020). Since polymerisation of ParA_F_ W362E-GFP was ATP-binding dependent, we next sought to determine if polymerisation required ATP hydrolysis and the conformational change to the ParA_F_-ATP* state. Different mutations in the same lysine residue (K120) have been reported to have different effects on ParA_P1_ ^(^Fung et al. 2001; Davis et al. 1996^;^ Vecchiarelli et al. 2013a). In ParA_F_, both K120R and K120Q have been shown to interact with ATP but fail to attain the ParA_F_-ATP* conformational state, but only ParA_F_ K120Q still retains its intrinsic ATP-hydrolysis activity similar to that of ParA_F_ (Ah-Senget al. 2013; Libante et al. 2001; Le Gall et al. 2016^)^. Therefore, we examined the effects of these ATP mutant variants (K120R and K120Q) on the polymerisation of ParA_F_ W362E-GFP. Both the double mutants, ParA_F_ K120R W362E and ParA_F_ K120Q W362E, failed to assemble into polymers and exhibited only diffuse cytoplasmic fluorescence, suggesting that a conformational change to ParA_F_-ATP* state but not ATP hydrolysis was necessary for the stabilisation of ParA_F_ filaments. We also substituted K120 with alanine in ParA_F_ W362E-GFP, and surprisingly, we observed that ParA_F_ K120A W362E-GFP was still able to assemble into polymers **(Fig. 7A)**. Furthermore, the percentage of cells having cytoplasmic filaments or the average filament length in the case of ParA_F_ K120A W362E-GFP were not significantly different than for ParA_F_ W362E-GFP **(Fig. 7 B and C)**. Thus, in contrast to the K120R and K120Q mutations, the K120A mutation did not prevent the polymerisation of ParA_F_ W362E-GFP.

The conformational change to the ParA_F_-ATP* state is essential for the nucleoid association and formation of ParA_F_ foci in the presence of the ParBS_F_ complex (Libante et al. 2001). Accordingly, ParA_F_ K120R and ParA_F_ K120Q are defective in foci formation (Hatano et al. 2007; Hu et al. 2021). We therefore reasoned that if ParA_F_ K120A retained the ability to undergo the conformational change to the ParA_F_-ATP* state, ParA_F_ K120A-GFP should assemble into foci like that of wild-type ParA_F_-GFP in the presence of ParBS_F_ complex **(Fig. 7D)**. Consistently, we observed that ParA_F_ K120A-GFP formed foci over the nucleoid region like ParA_F_-GFP. Quantification of the distribution of foci per cell showed only subtle differences between ParA_F_-GFP and ParA_F_ K120A-GFP **(Fig. 7E)**. Taken together, these results suggest that ParA_F_ K120A undergoes the conformational change to ParA_F_-ATP* state and that the assembly into cytoplasmic filaments by the ParA_F_ W362E-GFP requires the formation of ParA_F_-ATP dimers and the conformational change to its ParA_F_-ATP* state.

### Polar residues E375 in the last C-terminal helix and R320 in helix 14 play a role in polymerisation and plasmid maintenance function of ParA_F_

Recent cryo-electron microscopic studies of ParA2_Vc_ have revealed a polymeric structure of ParA bound to DNA, and the last C-terminal helix 16 constituted an important oligomer interface in the nucleoprotein filament complex (Parker et al. 2021). Moreover, we had earlier reported that the last C-terminal helix was essential for function in ParA_F_. In a study involving multiple point mutations of the charged residues in the C-terminal helix 16 of ParA_F_, we had found that E375A impacted nucleoid localisation of ParA_F_ *in vivo* (Mishra et al. 2022). Therefore, we introduced E375A into ParA_F_ W362E-GFP and tested their effects on the polymerisation of ParA_F_. Further, we carried out mutational studies in the conserved residues proximal (e.g. R320 and K329) to the previously identified nsDNA binding residues and tested their effects on ParA_F_ W362E-GFP. The double mutants, ParA_F_ W362E E375A-GFP and ParA_F_ W362E R320A-GFP, failed to assemble into polymers and exhibited diffuse fluorescence throughout the cell **(Fig. 8A)**, suggesting that E375 in the last C-terminal helix and R320 might be involved in the polymerisation of ParA_F_. Quantification of total fluorescence intensity in the cells expressing ParA_F_ W362E-GFP, ParA_F_ W362E E375A-GFP or ParA_F_ W362E R320A-GFP did not show any significant differences, suggesting that the protein levels were similar in these cells **(Fig. 8B)** and that the mutations R320A and E375A affected polymerisation of ParA_F_.

**Figure 8.**
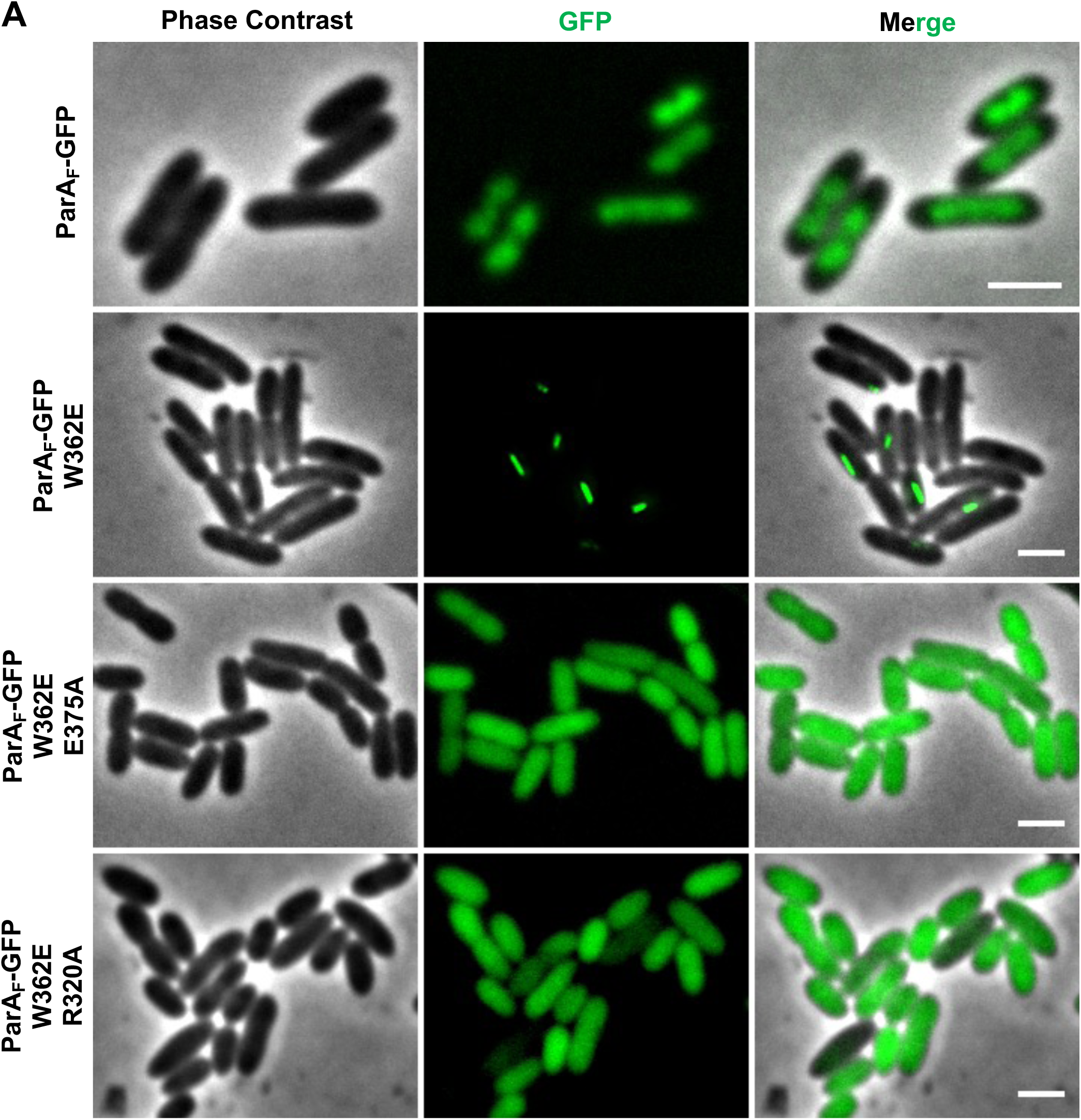
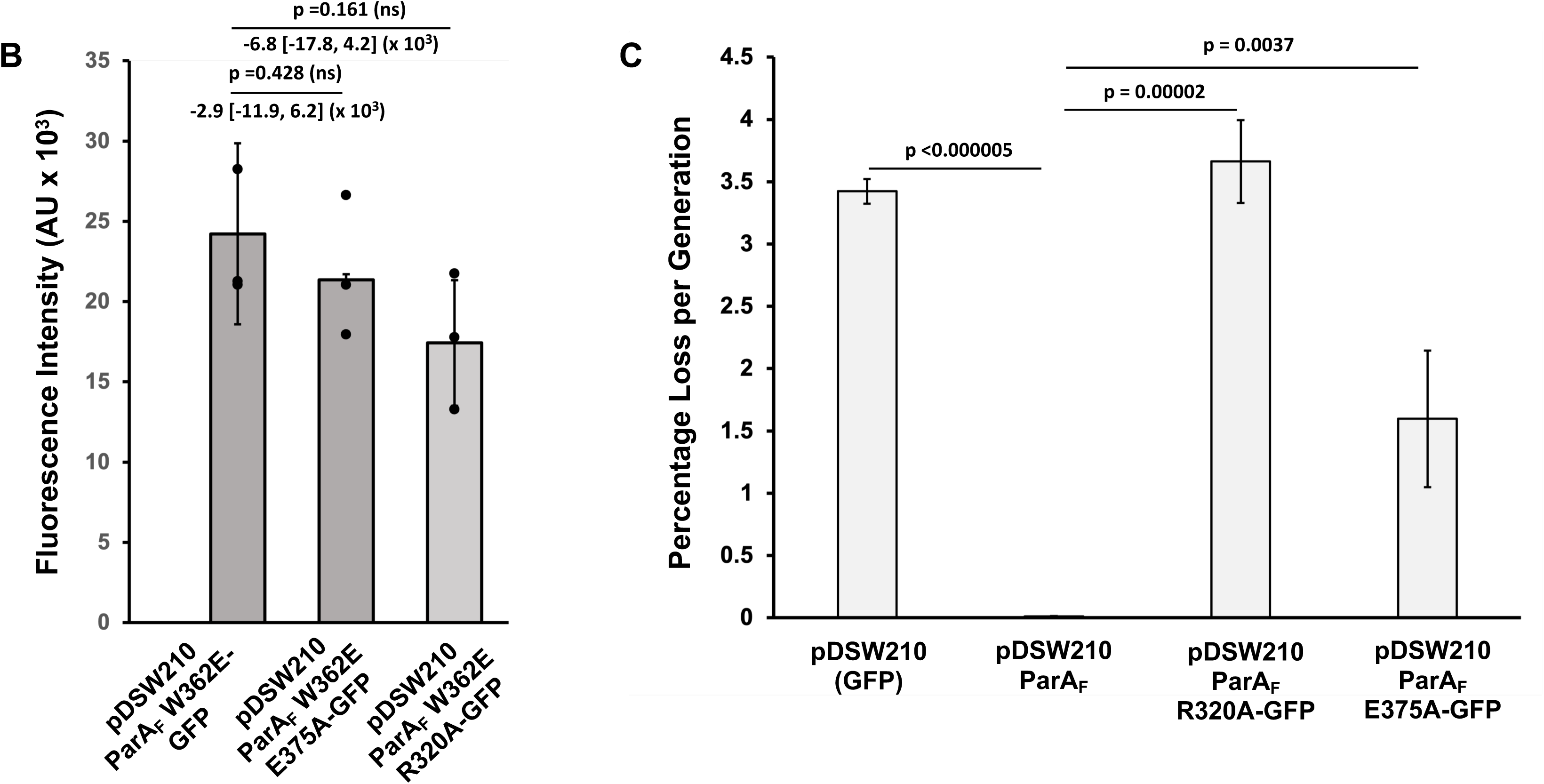
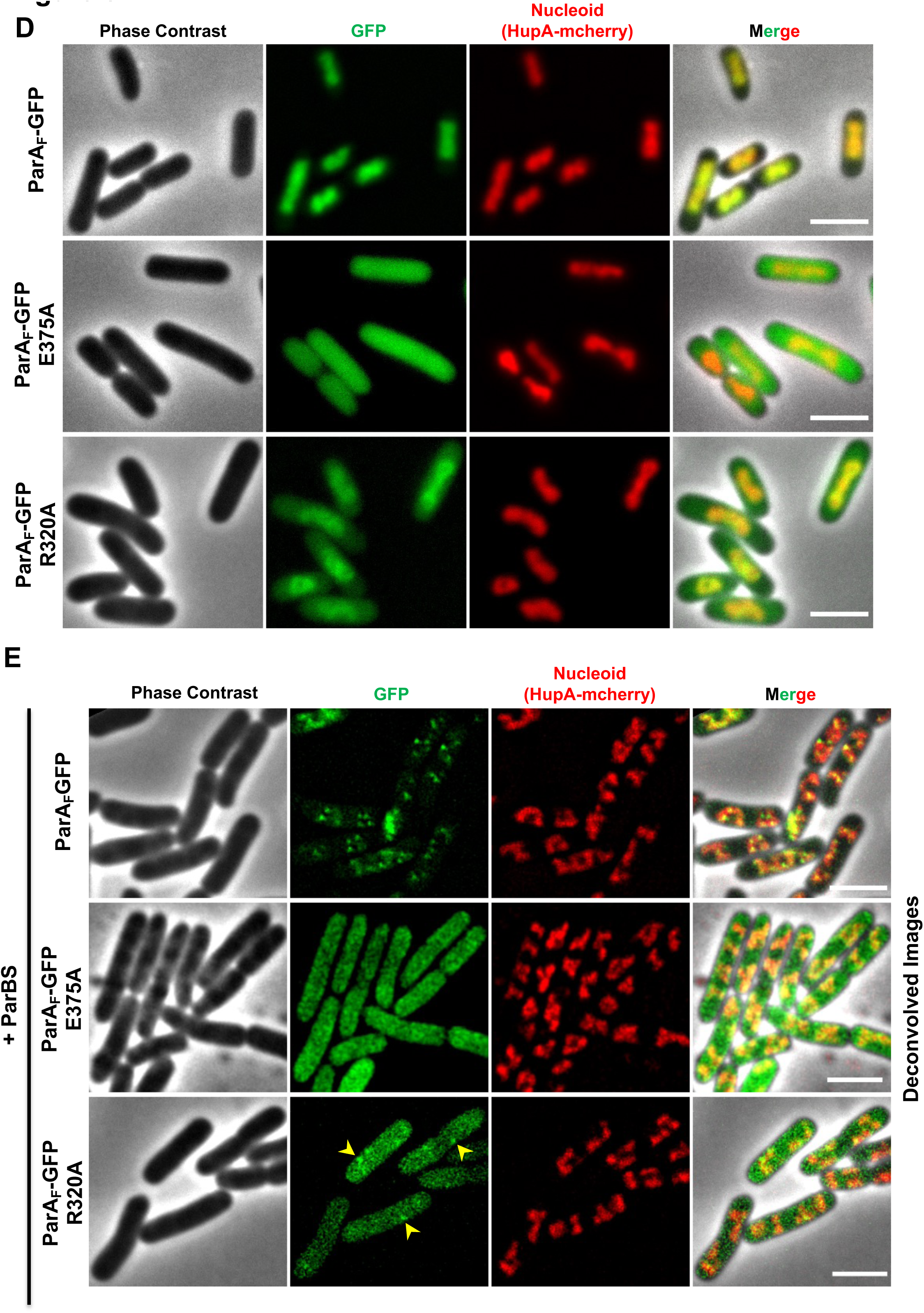
Identification of E375 in helix 16 and R320 in helix 14 as possible interface residues critical for polymerisation of ParA_F_. **(A)** While ParA_F_ W362E-GFP assembles into filaments, introducing either E375A or R320A abolishes polymerisation. **(B)** Quantification of the total fluorescence intensities (sum intensity) in cells expressing ParA_F_ W362E-GFP, ParA_F_ W362E E375A-GFP or ParA_F_ W362E R320A-GFP showing no significant differences in protein levels. Mean values from three independent biological replicates (N=3, number of cells counted was>400 for each replicate) are plotted. The values from the replicates are shown as black dots, and the mean value is plotted as a bar, with the SD values as the error bars. The p-values and the effect size are mentioned. The effect size is given as the difference in the means, and the 95% CI is indicated within parentheses. **(C)** Cultures expressing ParA_F_ W362E E375A-GFP or ParA_F_ W362E R320A-GFP fail to main the mini-F plasmids (*ΔparA*, *parBS*^+^) unlike cells expressing WT-ParA_F_-GFP. Error bars represent SD values. **(D)** Unlike ParA_F_-GFP, which co-localises with the bacterial nucleoid, ParA_F_ E375A-GFP and ParA_F_ R320A-GFP are distributed in the cytoplasm. However, ParA_F_ R320A-GFP shows mild localisation on the nucleoids compared to ParA_F_ E375A-GFP. **(E)** In the presence of the partitioning complex, ParA_F_ E375A-GFP fails to form foci and remains distributed throughout the cytoplasm, but ParA_F_ R320A-GFP shows weak foci formation. ParA_F_-GFP, as expected, appears as a haze or foci. An *E. coli* MG1655 derivative strain expressing carrying HupA-mCherry from its chromosomal locus was used to visualise the nucleoid, and cultures were induced with 100 µM IPTG for 2 hours to allow for the expression of ParA_F_-GFP, ParA_F_ E375A-GFP or ParA_F_ R320A-GFP from the pDSW210 vector. Scale bars represent 2 µm.

**Figure 9.**
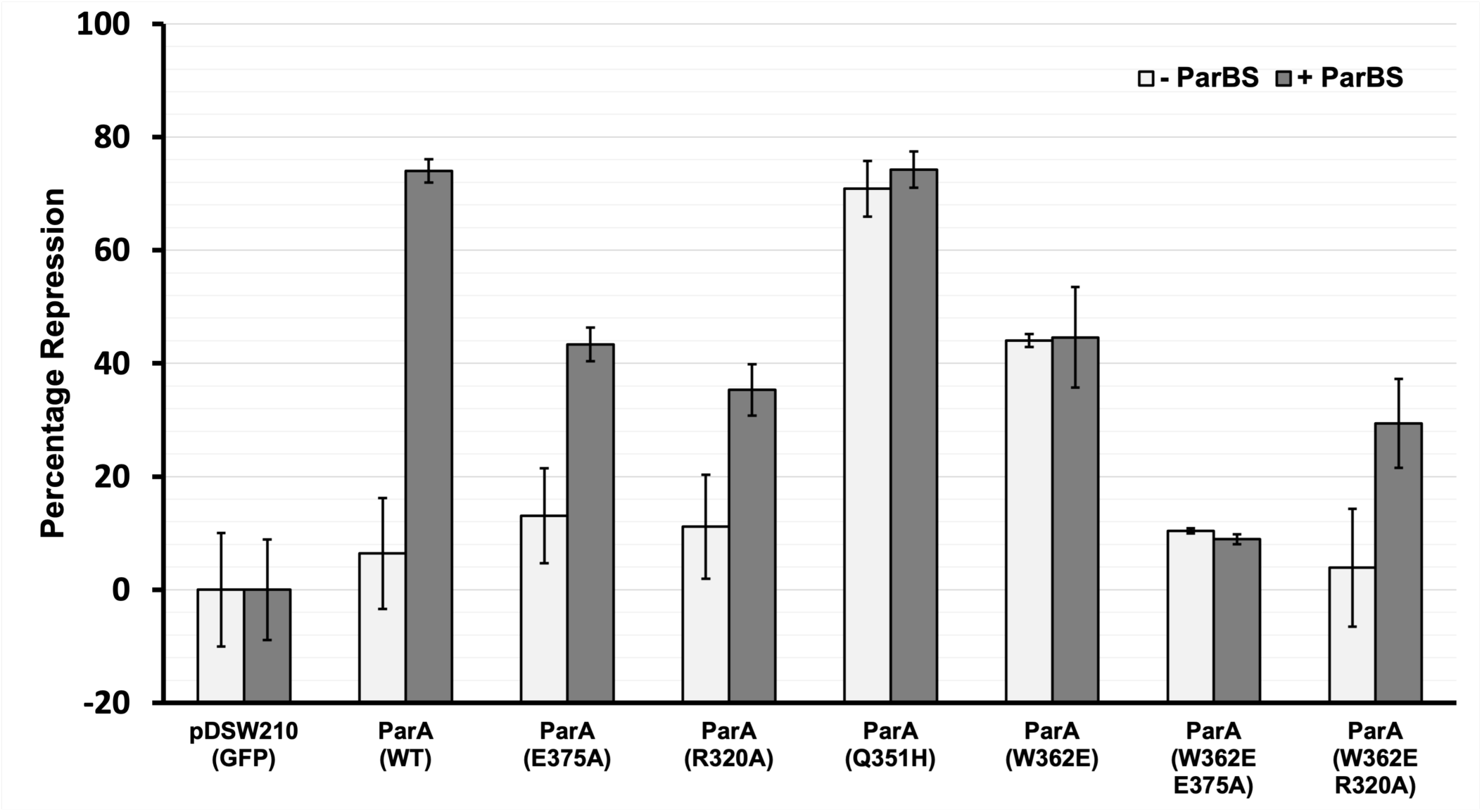
E375A and R320A mildly compromise the repressor activity of ParA_F_ but remain responsive to the ParBS_F_ partitioning complex. Quantitative β-galactosidase assay showing the repression activity by ParA_F_ and the various mutants on the P_parA_ promoter. While the repression of ParA_F_ is enhanced by the presence of the ParBS complex, ParA_F_ Q351H-GFP and ParA_F_ W362E-GFP strongly repress even in the absence of the ParB. Like ParA_F_, ParA_F_ E375A and ParA_F_ R320A show increased repression activity in the presence of the ParBS complex than in the absence of ParBS, albeit to a lesser extent compared to wild-type ParA_F_. The introduction of E375A or R320A mutations into ParA_F_ W362E also drastically reduces their ability to repress LacZ expression. Miller units were calculated as described in the materials and methods section, and the values obtained were normalised with respect to the vector control, pDSW210-GFP. Further, since the expression of LacZ in the vector control culture (in the absence of ParA_F_) was the maximum, normalised Miller units obtained in the case of pDSW210-GFP were considered as 100 % expressed or as 0 % repressed. Error bars represent the standard deviation of the mean (SD). Statistical significance test on the effect of ParBS on the repression activity was carried out using the estimated p-values from a one-tailed Student’s t-test (since repression values are only expected to be higher in the presence of ParBS) and are indicated in the graph.

Next, in order to ascertain if polymerisation and, thereby, residues E375 and R320 were important for ParA_F_ function, we determined if plasmid maintenance was affected by the E375A and R320A mutations. As described above, we utilised the two-plasmid system, wherein we assessed the stability of the mini-F plasmid (pDAG198; *ΔparA_F_, parBS^+^*) and expressed ParA_F_ or its variants from P*_trc_* promoter in the pDSW210 vector. We had earlier shown that ParA_F_ E375A-GFP exhibited mild plasmid loss compared to ParA_F_-GFP (Mishra et al. 2022). Consistently, we found that ParA_F_ E375A-GFP showed a plasmid loss of 1.6 ± 0.55 (SD, n = 3, at least 400 colonies were tested for each experiment) per generation **(Fig. 8C)**. ParA_F_ R320A-GFP showed a higher plasmid loss of 3.6 ± 0.33 (SD, n = 3, at least 400 colonies were tested for each experiment) and comparable to the plasmid loss in cultures lacking ParA_F_ [3.4 ± 0.1 (SD, n = 3, at least 400 colonies were tested for each experiment)] **(Fig. 8C)**. Further, we had also shown that ParA_F_ E375A-GFP did not bind DNA and failed to form foci in the presence of ParBS_F_ complex (Mishra et al. 2022). We thus tested if ParA_F_ R320A-GFP also behaved similarly and assessed its localisation in both the presence and absence of the ParBS_F_ complex. We used a strain of *E. coli* expressing HupA-mCherry to visualise the nucleoid. As a control, we also used ParA_F_-GFP, and as expected, it was found to be localised on the nucleoid in the absence of the ParBS_F_ complex. Consistent with our results, ParA_F_ E375A-GFP was found to be evenly distributed throughout the cell in the absence of the ParBS_F_ complex **(Fig. 8D)**. On the contrary, ParA_F_ R320A-GFP showed mild localisation to the nucleoid, but unlike ParA_F_-GFP, fluorescence was found to be distributed in nucleoid free regions of the cell as well **(Fig. 8D)**. Further, in the presence of the ParBS_F_ partitioning complex, only ParA_F_-GFP exhibited foci formation and both ParA_F_ E375A and ParA_F_ R320A resulted in predominantly diffuse fluorescence throughout the cytoplasm of the cell. However, ParA_F_ R320A exhibited mild foci consistent with its slight nucleoid association **(Fig. 8E)**. Our results here show that E375 and R320 are crucial for the polymerisation of ParA_F_, and mutations in these residues affect the nucleoid localisation and plasmid maintenance function of ParA_F_. These results further suggest that oligomerisation or formation of higher order structure by ParA_F_ might be crucial for plasmid partitioning *in vivo*.

### E375A and R320A mutations affect the super repression activity of ParA_F_ W362E

We next tested if the introduction of E375A or R320A into ParA_F_ W362E also abrogated their ability to act as super-repressors of transcription at the P*par_F_* promoter site using the above-described LacZ reporter and β-galactosidase assays.

Quantification of the β-galactosidase activity in the DLT1127 strain (P*_parA_*::*lacZ::kan*) carrying the various ParA_F_ constructs showed that the double mutants, ParA_F_ W362E E375A and ParA_F_ W362E R320A exhibited a decreased ability to repress LacZ expression as compared to ParA_F_ W362E, indicating that E375A and R320A mutations affected their ability to function as super-repressors **(Fig. 9)**. We used both null hypothesis statistical testing (NHST) and estimation statistics (Claridge-Chang and Assam 2016; Ho et al. 2019) to assess statistical significance of the data. The effect size, defined as the difference in the means, the precision of the difference given by the 95 % CI, and p-values for NHST are given in **Table 1 (i – iii)**. While the introduction of R320A into ParA_F_ W362E led to a difference of −40.1 [−57.8, −22.4] as compared to ParA_F_ W362E, the effect size in the case of ParA_F_ W362E E375A was −33.6 [−36.5, - 30.8] **(Table 1 i)**.

**Table 1.** Table showing the statistical analysis of the quantitative β-galactosidase assays for estimating promoter repression activity by the various ParA_F_ mutants. The table shows the statistical analysis for the repression activity by various ParA_F_ mutants in comparison to the vector control (pDSW210-GFP) or with ParA_F_, where the repression was normalised to 0 %. Estimation statistics were also used to assess the statistical significance of the repressor functions. Effect size is defined as the difference between the means of samples being compared, and the 95 % CI of the difference is given within the square parentheses. Statistical analysis showing the p-values and effect size, along with the 95 % CI of the difference, is tabulated. **(i)** Statistical analysis for the repression activity by ParA_F_ W362E E375A and ParA_F_ W362E R320A in comparison to ParA_F_ W362E showing the effect size along with the 95 % CI of the difference. The double mutants ParA_F_ W362E E375A and ParA_F_ W362E R320A show reduced repression of the P_parA_ promoter than ParA_F_ W362E with an effect size of −33.6 [−36.5, −30.8] and −40.1 [−57.8, −22.4], respectively. **(ii)** In the presence of ParBS, ParA_F_ E375A and ParA_F_ R320A show a reduced repression activity than the WT-ParA_F_ with an effect size of −30.7 [−43.7, −17.6] and −38.7 [−52.6, −24.8], respectively. **(iii)** Table shows the statistical analysis for the repression activity by various ParA_F_ mutants in comparison to the vector control (pDSW210-GFP). The effect size and the 95 % CI of the difference in the absence of ParBS as well as in the presence of ParBS are tabulated. In the absence of ParBS, ParA_F_ Q351H and ParA_F_ W362E act as super-repressors, showing an effect size of 70.8 [55.3, 86.4] and 44 [30, 58], respectively compared to pDSW210-GFP. When compared to ParA_F_, ParA_F_ Q351H and ParA_F_ W362E show an effect size of 64.4 [48.9, 80] and 37.6 [23.6, 51.6], respectively. The statistical significance of the results for the effect of the ParBS complex is also tabulated. Although ParA_F_ E375A and ParA_F_ R320A have reduced repression in the presence of ParBS compared to ParA_F_, they are responsive to ParB and their ability to repress is still enhanced by the presence of the ParBS complex. While ParA_F_ shows an effect size of 67.6 [51.5, 83.7], ParA_F_ E375A and ParA_F_ R320A show an effect size of 30.3 [16, 44.6] and 24.2 [7.7, 40.6], respectively. Results that are deemed statistically significant are shown in bold. P-values were obtained from a one-tailed Student’s t-test (since repression values are expected to be only higher (0 % or greater than experimental samples lacking ParBS). P-values ≥ 0.05 (after rounding off to 2 decimals) were considered statistically non-significant. Effect sizes are easier to interpret, with negative values indicating reduced repression ability and positive values indicating increased repression than the test sample. The 95 % CI of the difference, when not encompassing zero, indicates a statistically significant difference.

| ParA | Effect Size (ParA) | P-value (1-tailed) | Effect Size (ParA) | P-value (1-tailed) |
| --- | --- | --- | --- | --- |
|  | - ParBS |  | + ParBS |  |
| ParA W362E | NA | NA | NA | NA |
| ParA W362E E375A | -33.6 [-36.5, -30.8] | 0.00002 | -35.7 [-56.8, -14.6] | 0.00628 |
| ParA W362E R320A | -40.1 [-57.8, -22.4] | 0.00275 | -15.2 [-40, 9.6] | 0.07322 |

(ii)
| ParA | Effect Size (ParA) | P-value (1-tailed) | Effect Size (ParA) | P-value (1-tailed) |
| --- | --- | --- | --- | --- |
|  | - ParBS |  | + ParBS |  |
| ParA | NA | NA | NA | NA |
| ParA E375A | 6.7 [-11.5, 24.8] | 0.21140 | -30.7 [-43.7, -17.6] | 0.00006 |
| ParA R320A | 4.7 [-14.2, 23.6] | 0.28783 | -38.7 [-52.6, -24.8] | 0.00009 |

| ParA | Effect Size (pDSW210) | P-value (1-tailed) | Effect Size (ParA) | P-value (1-tailed) | Effect Size (pDSW210) | P-value (1-tailed) | Effect Size (ParA) | P-value (1-tailed) | Effect Size (ParBS) | P-value (1-tailed) |
| --- | --- | --- | --- | --- | --- | --- | --- | --- | --- | --- |
|  | - ParBS |  |  |  | + ParBS |  |  |  |  |  |
| pDSW210-GFP | NA | NA | -- | -- | NA | NA | -- | -- | NA | NA |
| ParA | 6.4 [-13.1, 25.9] | 0.23681 | NA | NA | 74 [61.3, 86.7] | 0.00007 | NA | NA | 67.6 [51.5, 83.7] | 0.00015 |
| ParA E375A | 13.1 [-5.1, 31.2] | 0.07939 | 6.7 [-11.5, 24.8] | 0.21140 | 43.4 [30.3, 56.4] | 0.00066 | -30.7 [-43.7, -17.6] | 0.00006 | 30.3 [16, 44.6] | 0.00209 |
| ParA R320A | 11.1 [-7.8, 30] | 0.11471 | 4.7 [-14.2, 23.6] | 0.28783 | 35.3 [21.5, 49.2] | 0.00180 | -38.7 [-52.6, -24.8] | 0.00009 | 24.2 [7.7, 40.6] | 0.00754 |
| ParA Q351H | 70.8 [55.3, 86.4] | 0.00020 | 64.4 [48.9, 80] | 0.00027 | 74.3 [61.1, 87.4] | 0.00009 | 0.2 [-12.9, 13.4] | 0.46074 | 3.4 [-6, 12.9] | 0.18623 |
| ParA W362E | 44 [30, 58] | 0.00082 | 37.6 [23.6, 51.6] | 0.00138 | 44.6 [27.1, 62.1] | 0.00178 | -29.4 [-46.9, -12] | 0.00250 | 0.6 [-13.8, 14.9] | 0.45868 |
| ParA W362E E375A | 10.4 [-13.4, 34.2] | 0.12910 | 4 [-19.3, 27.3] | 0.31173 | 8.9 [-12.3, 30.1] | 0.13652 | -65.1 [-70.2, -60.1] | 0.00002 | -1.5 [-4.4, 1.4] | 0.07948 |
| ParA W362E R320A | 3.9 [-25.6, 33.4] | 0.35112 | -2.5 [-31.6, 26.6] | 0.40120 | 29.4 [4.5, 54.3] | 0.01650 | -44.6 [-58.7, -30.6] | 0.00103 | 25.5 [-14.2, 65.2] | 0.05498 |

We also assessed the effect of the E375A and R320A single mutations on the repression ability of ParA_F_. In the absence of the ParBS_F_ complex, the β-galactosidase activity in the strain carrying ParA_F_ E375A or ParA_F_ R320A was not significantly different from those carrying ParA_F_ **(Table 1 ii)**. However, in the presence of the ParBS_F_ complex, both ParA_F_ E375A and ParA_F_ R320A repressed LacZ expression with a difference of 43.4 [30.3, 56.4] and 35.3 [21.5, 49.2], respectively, as compared to the vector control (pDSW210-GFP) **(Table 1 iii)**. On the contrary, ParA_F_ strongly reduced the expression of LacZ with an effect size of 74 [61.3, 86.7] compared to pDSW210-GFP. Thus, in comparison to ParA_F_, ParA_F_ E375A and ParA_F_ R320A showed a slightly reduced level of repression and had an effect size of −30.7 [−43.7, −17.6] and −38.7 [− 52.6, −24.8], respectively **(Table 1 ii)**., showing that ParA_F_ E375A and ParA_F_ R320A were moderately defective in repressing the transcriptional activity from the P*_parA_* promoter. Nonetheless, ParA_F_ E375A and ParA_F_ R320A showed enhanced transcriptional repression in the presence of the ParBS_F_ complex as compared to its absence, with an effect size of 30.3 [16.0, 44.6] and 24.2 [7.7, 40.6] **(Table 1 iii)**, suggesting that the E375A and R320A mutations did not affect the ParB and nucleotide binding functions of ParA_F_.

## Discussion

DNA partitioning by the ParA family of proteins, such as ParA_F_ (SopA), depends upon its ability to bind the nucleoid. It is the nucleoid-bound ParA_F_ that facilitates the process of plasmid segregation by interacting with the ParB-*parC* complex (SopB-*sopC*), which is essentially the plasmid cargo. While initial reports indicated that cytomotive filaments of ParA_F_ were responsible for the plasmid movements away from the mid-cell (Lim et al. 2005; Leonard et al. 2005; Gitai 2006; Hatano et al. 2007; Bouet et al. 2007), studies from several species of ParA proteins have shown that the ParA proteins interact as ATP-bound dimers with DNA (Vecchiarelli et al. 2010, 2013a; Havey et al. 2012; Lim et al. 2014; Zhang and Schumacher 2017; Köhler et al. 2022; Leonard et al. 2005; Hwang et al. 2013). However, recent studies have suggested that the binding of ParA to the nucleoid is cooperative and a cryo-EM study of ParA2_Vc_ revealed a filament structure of ParA bound to the DNA, suggesting that cooperativity could possibly involve oligomerisation of ParA (Chodha et al. 2023; Köhler et al. 2022; Parker et al. 2021). Thus, the exact function and mechanism of assembly of ParA into higher-order structures remains unclear. Polymerisation of ParA_F_ has been demonstrated *in vitro* (Bouet et al. 2007) in the presence of ATP, and filaments have also been observed in a few cells expressing wild-type ParA_F_ ^(^Lim et al. 2005; Hatano et al. 2007^)^.

A spontaneous mutant of ParA_F_ (M315I Q351H) has been shown to form static polymeric structures in *E. coli* (Lim et al. 2005). Our studies reported here show that another mutation at position W362 of ParA_F_ to either E or A also resulted in the assembly of ParA_F_ into cytoplasmic polymers similar to those formed by ParA_F_ M315I Q351H (SopA1). Further, a single mutation Q351H was sufficient to stabilise ParA_F_ filaments. Similar assembly of polymers by ParA_F_ M315I Q351H (SopA1), ParA_F_ Q351H, W362E or the W362A mutants in the absence of the ParBS_F_ complex or in a heterologous eukaryotic host, fission yeast, suggested that the mutations helped stabilise an otherwise transient polymeric form of ParA_F_ independent of the partition complex. Moreover, both ParA_F_ Q351H or ParA_F_ W362E were able to induce the polymerisation of wild-type ParA_F_ when co-expressed and resulted in the formation of cytoplasmic filaments of WT-ParA_F_-GFP, even though ParA_F_ Q351H exhibited a much stronger effect compared to ParA_F_ W362E.

However, both the mutants (Q351H and W362E), like SopA1 (M315I Q351H), were defective in plasmid maintenance, showing that the static or stable polymers of ParA_F_ did not drive DNA segregation. Further, super-resolution imaging using 3D-SIM revealed that ParA_F_ Q351H-GFP and ParA_F_ W362E-GFP filaments exhibited nucleoid binding defects, consistent with their inability to maintain plasmids. There is ample evidence supporting non-specific DNA binding and dynamic localisation of ParA_F_ to the nucleoid driving the process of plasmid segregation. These results show that polymerisation in the absence of nucleoid binding does not facilitate plasmid partitioning and maintenance. Moreover, *in vitro* polymerisation studies using light-scattering in the presence of nsDNA have implicated DNA as a polymerisation inhibitor (Bouet et al.2007). Thus, in contrast to Soj and ParA2_Vc_ that form nucleoprotein filaments in the presence of non-specific DNA (Leonard et al. 2005; Hui et al. 2010; Parker et al. 2021), the lack of non-specific DNA binding by ParA_F_ M315I Q351H (SopA1), ParA_F_ Q351H and ParA_F_ W362E could have possibly induced polymerisation. However, K340A, a mutation known to abolish the nsDNA binding in ParA_F_, does not assemble into such micron-long filaments (Castaing et al. 2008), suggesting that the residues Q351 and W362 in the C-terminal region may play a crucial stabilising role in polymerisation of ParA_F_.

Furthermore, the polymers formed by ParA_F_ Q351H or ParA_F_ W362E underwent depolymerisation within 20 minutes when new protein synthesis was stopped, suggesting a need for a critical protein concentration to be maintained in cells for polymerisation. Disassembly of filaments upon inhibition of protein synthesis suggested that the maintenance of the polymers required continual protein synthesis and the existence of a cytoplasmic pool of ParA_F_ Q351H or ParA_F_ W362E proteins as well as the polymeric form. Consistent with the inability of ParA_F_ Q351H or ParA_F_ W362E proteins to associate with the bacterial nucleoid, both mutations resulted in super-repression of transcription from the ParA_F_ promoter (P*_parA_*), i.e. enhanced transcriptional repression in the absence of the ParBS_F_ complex. These findings are consistent with the recent reports on ParA_P1_ DNA binding mutants. The absence of nsDNA binding results in an excess-free pool of ParA that becomes available to bind the promoter and function as super-repressors (Baxter et al. 2020). Thus, the inability of ParA_F_ Q351H and ParA_F_ W362E to bind nsDNA might create an excess of the free cytoplasmic pool of the protein, which can then exhibit heightened auto-repression activity. We believe that while the filaments may not associate with the nucleoid, the available cytoplasmic pool might interact with the specific ParA_F_ operator sites, forming a stable complex, acting to prevent transcription effectively. Thus, the mutations (Q351H and W362E) do not impair the ability of ParA_F_ to bind with specific DNA. These results suggested that the mutations Q351H and W362E did not affect the overall structural integrity of the protein. The mutations Q351H and W362E possibly stabilise the trimer of dimer structures of ParA_F_ that have been described recently (Boudsocq et al. 2021), enabling the formation of a tighter complex with the ParA_F_ promoter sites and might be conducive to the organisation of larger filamentous structure beyond a threshold of protein concentrations.

Consistent with the earlier *in vitro* studies (Fung et al. 2001; Libante et al. 2001), polymerisation of ParA_F_ Q351H and W362E was an ATP-dependent process since introducing a mutation that prevents ATP-binding (K120E) completely abolished filament assembly. Mutational studies of the conserved K120 residue in the Walker A motif have also shown that the residue is also crucial for the necessary conformational change from ParA_F_-ATP to ParA_F_-ATP* (Libante et al. 2001; Ah-Seng et al. 2013). Accordingly, earlier localisation studies on the various ParA_F_ K120 substitutions have shown that the K120R and K120Q mutants fail to form foci in the presence of the ParBS_F_ complex (Hatano et al. 2007). The ability of ParA_F_ K120A to form foci similar to that of wild-type ParA_F_ in the presence of the ParBS_F_ complex suggests that among the various K120 mutants, K120A alone could support the conformational change from ParA_F_-ATP to ParA_F_-ATP*. Further, our studies on combining the K120 mutations with W362E show that only ParA_F_ K120A W362E retained the ability to assemble into filaments. Moreover, previous *in vitro* studies had indicated that ParA_F_ assembly into filaments possibly depended upon the conformational change induced by ATP. These results suggest that, like nsDNA binding, polymerisation of ParA_F_ was also dependent upon the conformational change to the ParA_F_-ATP**\*** state. It is plausible that the mutations Q351H and W362E abolish the ATP hydrolysis activity of ParA_F_ and stabilise the polymeric state. Interestingly, while the ATP-hydrolysis mutants K120Q or K120A by themselves do not form ParA filaments, yet another mutant, ParA_F_ V125A, which is also defective in ATP-hydrolysis, has been shown to assemble into polymers *in vivo* (Castaing 2009). Thus, inhibition of ATP-hydrolysis by mutations in the residues other than the conserved K120 in the Walker A motif might result in the stabilisation of ParA_F_ polymers.

Clearly, further biochemical experiments on these mutants will be required in the future to address these possibilities and gain insights. On the other hand, we have utilised the ability of ParA_F_ W362E to form stable filaments to identify new residues crucial for polymerisation for ParA_F_. Mutations in residues proximal to the previously identified nsDNA binding residues (Castaing et al. 2008) and the C-terminal helix ^(^Mishra et al. 2022^)^ in ParA_F_ revealed R320 (in helix 14) and E375 in helix 16, respectively, as being critical for the polymerisation of ParA_F_ W362E. Both ParA_F_ W362E R320A and ParA_F_ W362E E375A failed to form filaments and exhibited reduced transcriptional repression activity at the ParA_F_ promoter, suggesting a role for E375 and R320 in oligomerisation/ polymerisation. Accordingly, the single mutants ParA_F_ R320A and ParA_F_ E375A also showed reduced promoter repression in the presence of the ParBS_F_ complex as compared to the WT-ParA_F_. Nonetheless, both ParA_F_ R320A and ParA_F_ E375A responded to ParB and were able to repress the transcriptional activity at the ParA_F_ promoter in the presence of the ParBS_F_ partitioning complex, albeit to a lesser extent than wild-type ParA_F_. Transcriptional repression at the ParA promoter requires nucleotide binding (Lemonnier et al. 2000; Libante et al. 2001; Ah-Seng et al. 2013; Baxter et al. 2020). These results, therefore, suggest that the mutations R320A and E375A retained their ability to bind nucleotide and associate with specific promoter DNA sequences to prevent transcription of ParA_F_ in response to ParBS_F_. Therefore, we infer that the loss of plasmid maintenance observed for ParA_F_ R320A and ParA_F_ E375A was not due to the lack of ATP binding by the mutants but possibly due to their defects in oligomerisation. These results are thus consistent with the recently proposed role of helix 14 and the last C-terminal helix 16 in intra-subunit contacts within ParA_Vc_ filaments (Parker et al., 2021) and co-operative binding of ParA to nsDNA, which may involve polymerisation, albeit transient (Parker et al. 2021; Chodha et al. 2023).

We envisage a model wherein upon ATP binding, ParA_F_ undergoes the conformational change to the ParA_F_-ATP**\*** state as proposed earlier and binds co-operatively to nsDNA within the high-density regions (Le Gall et al. 2016; McLeod et al. 2017) via assembly into higher-order structures (Hui et al. 2010; Parker et al. 2021). However, the stabilisation of oligomeric structures by the mutants Q351H or W362E results in the formation of long cytoplasmic filaments incompatible with nsDNA binding, resulting in a failure to associate with the nucleoid. Although further biochemical and electron microscopy studies are required to elucidate the oligomerisation of ParA_F_ and its precise roles *in vivo*, one possibility could be that the polymerisation of ParA_F_ could act as a sequestration mechanism to regulate any excess ParA_F_ from the system. Alternately, our studies here lead us to speculate that the last C-terminal helix of ParA_F_ might be a key player in facilitating nsDNA binding and the segregation of plasmids by regulating ParA_F_ oligomerisation **(Fig. 10)**. Further identification of ParA mutants specifically defective in polymerisation but not in ATP-binding and formation of the ATP-bound sandwich dimers or ParA-ATP* state, and similar studies across other members of the ParA family of DNA partitioning proteins and their genetic and biochemical characterisation should lead to a better insight into the physiological role of polymerisation by the ParA family of proteins.

**Figure 10.**
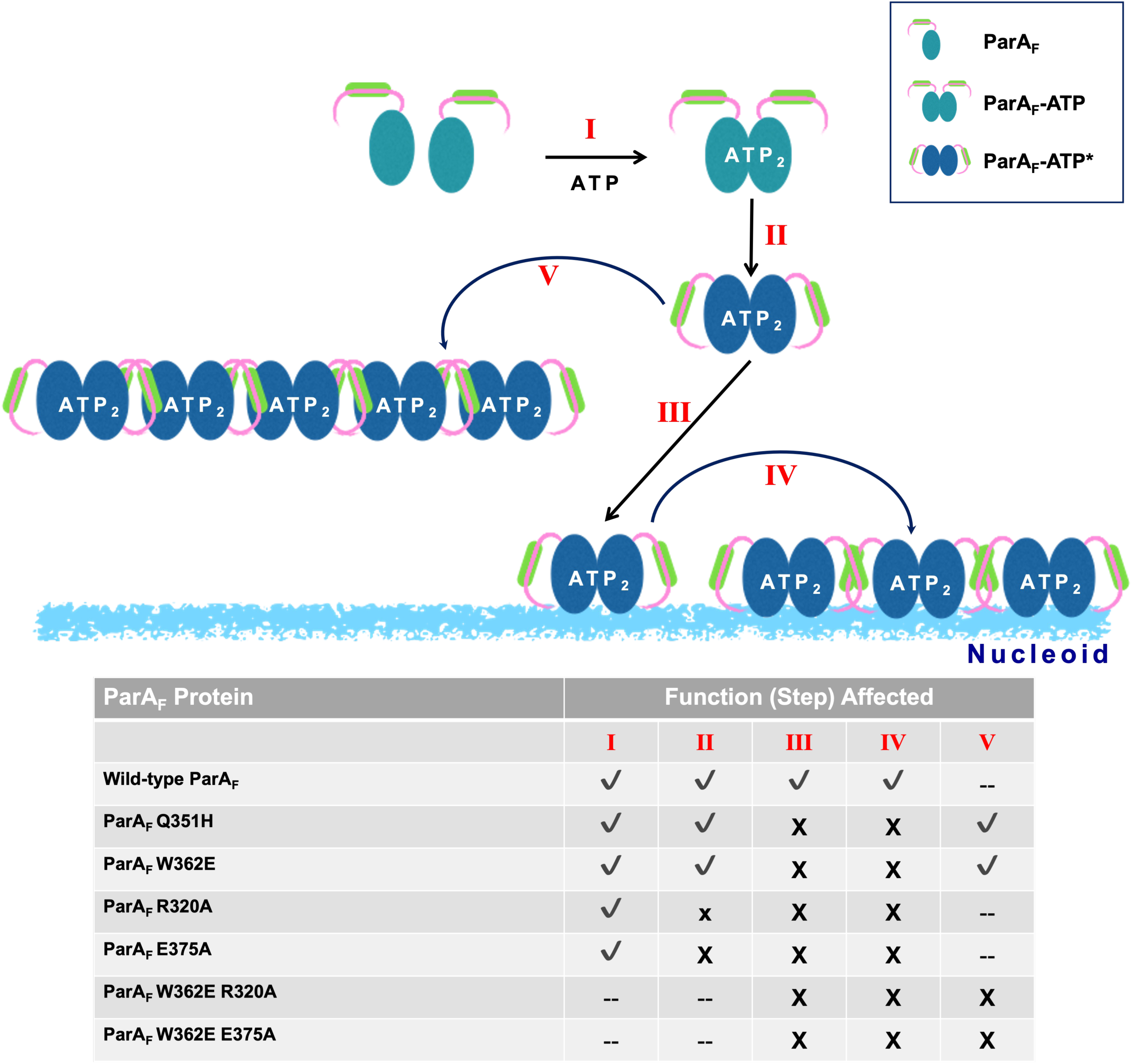
Schematic showing the effect of various mutations on the oligomerisation of ParA_F_ protein. Monomeric apo-ParA_F_ binds ATP to form the ATP sandwich dimer and undergoes a conformational change, resulting in the ParA_F_-ATP* state. The ParA_F_-ATP* binds the bacterial nucleoid as an ATP sandwich dimer, which could further undergo cooperative binding involving transient oligomerisation. The various steps are marked in roman numerals and the effect of various mutations is tabulated. A tick mark (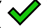) shows activity, a small cross mark shows (x) partial impairment of activity and a cross mark (X) shows the step likely affected by the mutation. The mutations Q351H and W362E result in the formation of stable cytoplasmic filaments that are incompatible with nsDNA binding (nucleoid localisation). The mutations in the C-terminal helix-14 (R320A) or helix-16 (E375A) abolish oligomerisation. The mutations R320A and E375A might possibly inhibit polymerisation either by blocking direct subunit interactions or preventing the conformational change to the ParA_F_-ATP* state.

## Materials and Methods

### Strains, Plasmids, Media and Growth Conditions

*E. coli* and *S. pombe* strains were cultured as described previously (Srinivasan et al. 2007; Mishra et al. 2021). While *E. coli* was grown at 37°C in LB medium, *S. pombe* was grown at 30°C in Yeast Extract with Supplements (YES) medium or Edinburgh Minimal Medium (EMM) supplemented with adenine (0.225 mg/ml), histidine (0.225 mg/ml) and leucine (0.225 mg/ml) or uracil (0.1125 mg/ ml) as necessary but lacking thiamine. Antibiotics carbenicillin and chloramphenicol in LB medium were used at a concentration of 100 μg/ mL and 34 μg/ mL, respectively. Kanamycin was used at 50 μg/ mL (for plasmids) or 25 μg/ mL in the case of single-copy chromosomal integrations. Chloramphenicol was added at a 50 µg/ mL concentration to inhibit protein synthesis in cells and at 50 µg/ mL concentration to condense nucleoids (Sun and Margolin 2004; Zusman et al. 1973). Cephalexin at a final concentration of 25 µg/ mL (Pogliano et al. 1997) was added to inhibit cell division for 30 minutes. *E. coli* DH5α was used for cloning purposes. All the plasmids used in this study were created using Gibson assembly (Gibson et al. 2009) or Restriction Free cloning (van den Ent and Löwe 2006) and site-directed mutations were introduced by the method of Stratagene QuikChange^TM^ using Q5 polymerase (NEB). In experiments relating to promoter repression, a stop codon (TAA) was introduced between the ParA_F_ and the GFP coding sequences, as previously reported (Mishra et al. 2022). The strains and plasmids used in this study have been listed in **Supplementary Table S1**, and the oligonucleotides used for generating the constructs have been listed in **Supplementary Table S2**. Expression of ParA_F_-GFP from the weak P_Trc_ promoter of the pDSW210 vector was achieved by adding 50 µM or 100 μM isopropyl-β-D-thiogalactopyranoside (IPTG) as mentioned in the text.

### Live Cell Imaging-

Live-cell imaging of *E. coli* and *S. pombe* was carried out as previously described (Srinivasan et al. 2007, 2008; Mishra et al. 2022) using an epifluorescence microscope (DeltaVision Elite^TM^) equipped with 7-color solid-state illumination (Spectra Light Engine from Lumencor^TM^). A 100 x oil immersion of NA 1.4 (UPLSAPO100XO) or a phase objective lens (PLN100XOPH) of NA 1.4 was used, and images were acquired using a CCD camera (CoolSNAP^TM^ HQ2). Excitation filters and emission filters of −390/18 and 435/48 (DAPI), 475/28 nm and 525/48 nm (GFP) and 575/25 nm and 625/45 nm (mCherry) were used, respectively. Briefly, cells were pelleted down by centrifuging at 2000 x g for 5 min and spotted on LB (for *E. coli*) or EMM without thiamine (for *S. pombe*) agarose pads (1.5 % agarose). Nucleoids in *E. coli* were visualised by incubating cells with 0.5 μg/ml 4′,6-diamidino-2-phenylindole (DAPI) for 15–20 min prior to centrifugation. Alternatively, an MG1655-derived strain of *E. coli* expressing HupA-mCherry from its chromosomal locus was used to visualise the nucleoid. Membranes were stained with FM4-64 and additives such as antibiotics, IPTG, 2% glucose, chloramphenicol at 50 ug/ml to prevent protein synthesis or 100 ug/ml to condense nucleoids (Zusman et al. 1973; Sun and Margolin 2004) and cephalexin at 25 µg/ml to prevent cell division (Pogliano et al. 1997) were all added to the agarose pads where mentioned in the text. 3D-SIM imaging was carried out using a DeltaVision OMX-SR Blaze microscope equipped with a PCO Edge 4.2 sCMOS camera, and a 60x, NA 1.42 oil-immersion objective lens was used to acquire raw images. An exposure time of 2 – 5 ms was used to acquire each image, and Z-stacks were obtained at a step size of 0.125 μm to cover the width of the bacterial cell. 3D-SIM images were reconstructed using the SI reconstruction module of SoftWorx^TM^ software. All images were acquired using SoftWorx^TM^ software and processed with Fiji v-1.53c or v-1.54f (Schindelin et al. 2012). The deconvolution of images was performed using SoftWorx^TM^ software’s in-built algorithm - DECON3D: 3D iterative constrained deconvolution with not more than 10 iterations.

### Plasmid Stability Assay

Plasmid stability assay was performed using a two-plasmid system (Ah-Seng et al. 2013), as described previously (Mishra et al. 2021, 2022). Briefly, bacterial cultures carrying the plasmids were sub-cultured 1:1000, maintained in exponential phase by sub-culturing (1:100) twice into a fresh LB media and allowed to grow for 40 generations. Various dilutions of the culture were plated on carbenicillin plates and incubated at 37 °C. Colonies were then patched on plates containing chloramphenicol to estimate the retention of the mini-F plasmid. The plasmid loss rate was calculated following the equation 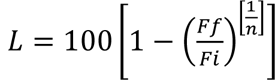 as reported previously (Ravin and Lane 1999). “Fi” is the initial fraction of cells containing plasmid, and “Ff” is the number of cells carrying the plasmid after 40 generations (n) of growth.

### Promoter repression assay

Qualitative assessment to detect the LacZ activity was carried out on LB agar plates containing IPTG (200 μM) and X-gal (40 μg/ mL). Overnight cultures carrying the various ParA_F_ constructs were diluted 1:100 into fresh LB, and growth was continued until the cultures reached an OD_600_ of 0.2. The cultures were induced with 100 μM IPTG for 2 hours. Subsequently, serial dilutions of the cultures (adjusted to the same OD_600_) were spotted on indicator plates-X-gal (40 μg/ mL) + IPTG (0.2 mM). These plates were then incubated overnight at 37 °C, following which they were imaged using a CCD colour camera on a mobile phone device and the white balance was corrected using the ImageJ plugin. For quantitative estimates of the LacZ expression, β-galactosidase assays were performed as described (Griffith and Wolf 2002). Cultures were centrifuged at 1523 x g, 10 min, 4 °C and the pellet was resuspended in Z-buffer (0.06 M Na_2_HPO_4_, 0.04 M NaH_2_PO_4_, 0.01 M KCl, 0.001 M MgSO_4_, pH 7.0). 50 μM β-mercaptoethanol was added to the Z-buffer immediately before use. A 2 mL culture adjusted to an OD_600_ of 0.6 was used for all assays. Cells were permeabilised by the addition of 20 μL Chloroform and 20 μL of 0.1 % SDS solution. Samples were then vigorously vortexed for one to 2 minutes and incubated at 28 °C for 10 minutes. Nitrophenyl-β-D-galactopyranoside (ONPG) at a final concentration of 0.13 mM was used as the substrate for the reaction and recorded as time point zero (*t_0_*) for the assay. When ONPG is hydrolysed by β-galactosidase into galactose and *o*-nitrophenol, the solution colour turns yellow. When sufficient yellow colour had developed (assessed visually), the reaction was stopped by the addition of 100 μL of 1 M solution of Na_2_CO_3_ and the time was recorded (in minutes). Values of the β-galactosidase activity were determined in Miller units using the formula: 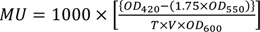, where MU is Miller Units, OD_420_, OD_550_ and OD_600_ are measure optical density at 420 nm, 550 nm and 600 nm, respectively. T is the total reaction time in minutes, and V is the volume of culture used in milli-litres (mL).

### Statistical Methods

A single colony of bacteria or a patch of yeast cells from freshly streaked out cells on agar plates from frozen glycerol stocks and grown independently were considered biological replicates (n). Except where mentioned in the text, every experiment was performed at least 3 or more times (independent biological replicates, n ≥ 3). All error bars represent the standard deviation from the central tendency value mean. In the case of the plasmid stability assays, the loss rate of mini-F plasmid was compared to the cultures that carried the wild-type ParA_F_, wherein the plasmid loss rate was always zero. Therefore, a one-sample t-test was used to calculate p-values for evaluating the statistical significance of the plasmid loss rates among various mutants. The online web server https://www.graphpad.com/quickcalcs/oneSampleT1/ was used to obtain the p-values.

For other experiments pertaining to the promoter repression assays, where p-values are mentioned, either two-tailed Student’s t-test or one-tailed Student’s t-test was used, wherein the repression values were already expected to be only higher, as in the case of vector control (normalised to zero) and ParBS (which is expected to enhance repression). As an alternative to null-hypothesis significance tests (NHST), we also used estimation statistics to determine the effect size and the precision of the effect size was given by the 95 % confidence intervals (Claridge-Chang and Assam 2016; Ho et al. 2019). Superplots were plotted as described (Lord et al. 2020) using Excel Office 365. Effect size is defined as the difference between the mean values of the samples being compared, and statistical significance is assumed when 95% CI values do not encompass zero (Cumming et al. 2007; Vaux 2014; Pollard et al. 2019).

## Acknowledgement

The authors thank all the members of the RS lab for their helpful suggestions and technical help. DM and NM received financial support and fellowship from DAE, which is acknowledged. RS is the recipient of research grants from DST-SERB (CRG/2021/000337) that supported this work. Intra-mural funding from the Department of Atomic Energy (DAE) is acknowledged. The authors sincerely thank Drs. Anjana Badrinarayanan (NCBS, TIFR, DAE) and Gayathri Pananghat for their suggestions, helpful discussions and comments on the manuscript. The authors thank Drs. Jean Yves-Bouet (LMGM, Toulouse, France), David Lane (CBI-Toulouse, France), Tushar Beuria (ILS, Bhubaneswar) and Mohan Chandra Joshi (JMI, New Delhi), for the generous gifts of strains and plasmid. pDSW210 was originally a gift from Dr. David Weiss (Univ. of Iowa), and the *E. coli ΔminB* strain was originally obtained from Dr. Lutkenhaus (Univ. of Kansas).

## Author Contributions

NM, DM and IAP performed experiments. NM and DM carried out microscopy, acquired images and analysed them. IAP performed and analysed certain experiments pertaining to promoter repression and analysed images. NM quantified, analysed and organised the data. NM and DM contributed to writing the draft manuscript. RS and NM edited the manuscript. RS conceptualised the work, RS and NM designed experiments, analysed and interpreted the data.

