## Supplementary Data for "Mutational analysis of the F plasmid partitioning protein ParA reveals novel residues required for oligomerisation and plasmid maintenance"

### Supplementary Figures

#### Supplementary Figure 1.

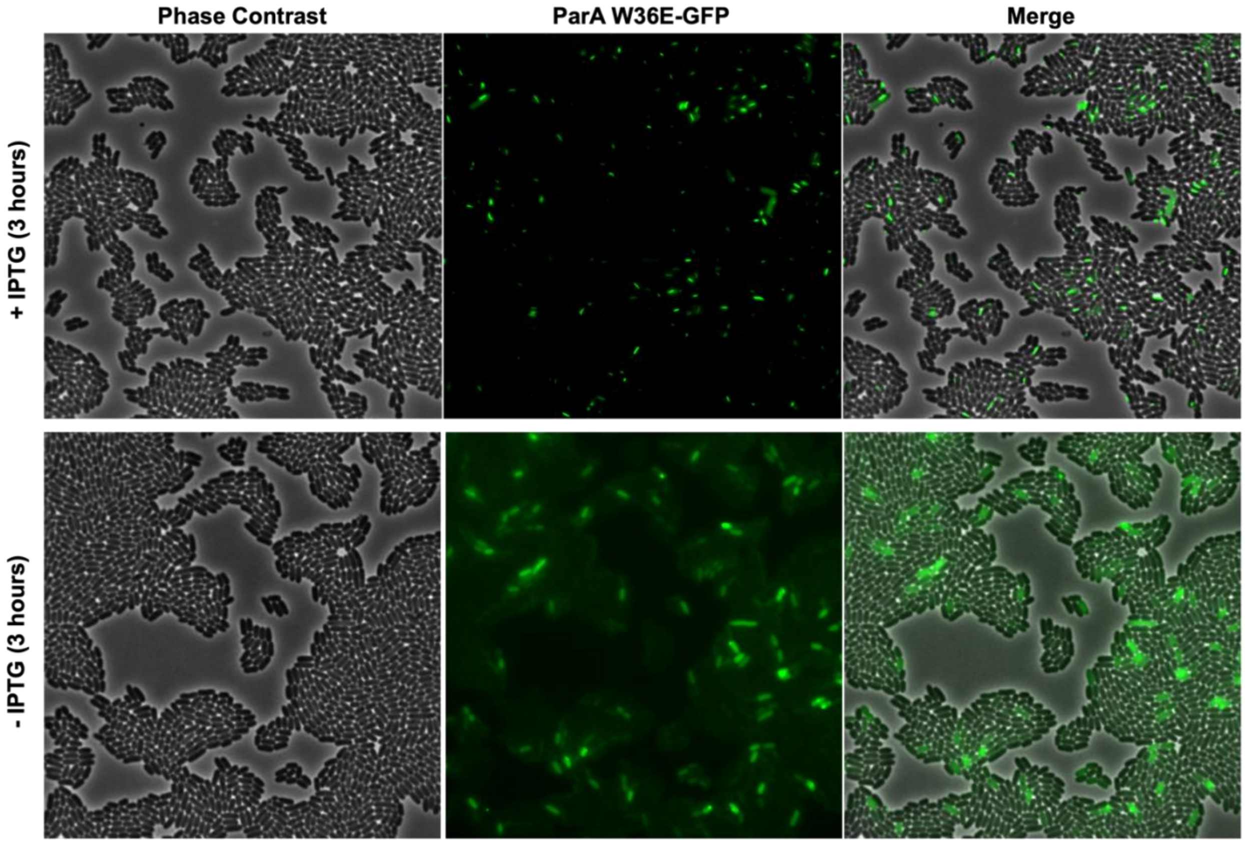


#### Supplementary Figure Legends

**Supplementary Figure 1. Disassembly of ParA$_F$ filaments assembled by W362E mutant in the absence of continued protein synthesis.**

ParA$_F$ W362E-GFP filaments disassemble in the absence of the inducer IPTG. *E. coli* MC4100 cells carrying pDSW210-ParA$_F$ W362E-GFP were induced 100 µM IPTG and then placed on LB-agarose slides with 100 µM IPTG **(top panel)** or without IPTG **(bottom panel)**.

### Supplementary Tables

#### Supplementary Table S1.

#### List of strains and plasmids used in the study.

| Strains | Name | Genotype | Source | Reference |
|---|---|---|---|---|
| **Bacterial strains** | | | | |
| | Source Name | | | |
| CCD219 | DH5α | *fhuA2 lac(del)U169 phoA glnV44 Φ80' lacZ(del)M15 gyrA96 recA1 relA1 endA1 thi-1 hsdR17* | Dr. Tushar K Beuria | Meselson and Yuan, 1968; Hanahan, 1985 |
| CCD253 | MC4100 | $F^-$ [*araD139*]B/r Δ(*argFlac*)169* $\lambda^-$ $e14^-$ *flhD5301* Δ(*fruK-yeiR*)725 (*fruA25*) *relA1 rpsL150* (strR) *rbsR22* Δ(*fimB- fimE*) 632(*::IS1*) *deoC1* | Dr. Tushar K Beuria | Casadaban, 1976 |
| CCD322 | | MG1655 *hupA-mCherry::kanR* | Dr. Mohan Chandra Joshi | Marceau et al., 2011; Fisher et al., 2013 |
| CCD358 | DLT1127 | *araD139* Δ(*ara leu*)7679, Δ*lacX74 galU galK rpsL thi hsdR2 mcrB* λRS88-kan-$P_{ParA_F}$::*lacZ* | Dr. David Lane | Ravin and Lane, 1999 |
| **Yeast Strains** | | | | |
| CCDY346 | MBY192 | ura4-D18, leu1-32, h- | Dr. Mohan Balasubramanian | Lab Collection |
| **Yeast Strains with plasmids** | | | | |
| CCDY58 | | MBY192/ pREP42-ParA$_F$-GFP | | This work |
| CCDY75 | | MBY192/ pREP42-ParA$_F$ M315I Q351H-GFP (SopA1-GFP) | Dr. Mohan Balasubramanian | This work |
| CCDY421 | | MBY192/ pREP42-ParA$_F$ Q351H-GFP | | This work |
| CCDY422 | | MBY192/ pREP42-ParA$_F$ W362E-GFP | | This work |

| **Plasmids** | | | |
|---|---|---|---|
| **Plasmid** | **Description** | **Vector backbone** | **Reference** |
| pCCD479 | GFP under weak $P_{trc}$ promoter | pDSW210 | Weiss et al., 1999 |
| pCCD494 | $ParA_F$-GFP | pDSW210 | Lim et al., 2005 |
| pCCD495 | $ParA_F$ M315I Q351H-GFP (SopA1-GFP) | pDSW210 | Lim et al., 2005 |
| pCCD960 | $ParA_F$ W362A-GFP | pDSW210 | This work |
| pCCD593 | $ParA_F$ Q351H-GFP | pDSW210 | This work |
| pCCD589 | $ParA_F$ W362E-GFP | pDSW210 | This work |
| pCCD51 | pREP42-GFP | pREP42 | Srinivasan et al., 2008 |
| pCCD83 | pREP42- $ParA_F$-GFP | pREP42 | This work |
| pCCD898 | $ParA_F$ Q351H-GFP | pREP42 | This work |
| pCCD899 | $ParA_F$ W362E-GFP | pREP42 | This work |
| pCCD810 | $ParA_F$-Stop-GFP | pDSW210 | This work |
| pCCD809 | $ParA_F$ W362E-Stop-GFP | pDSW210 | This work |
| pCCD825 | $ParA_F$ Q351H-Stop-GFP | pDSW210 | This work |
| pCCD687 | $ParA_F$ W362E K120E-GFP | pDSW210 | This work |
| pCCD682 | $ParA_F$ W362E K120Q-GFP | pDSW210 | This work |
| pCCD879 | $ParA_F$ W362E K120R-GFP | pDSW210 | This work |
| pCCD677 | $ParA_F$ W362E K120A-GFP | pDSW210 | This work |
| pCCD498 | $ParA_F$ K120A-GFP | pDSW210 | This work |
| pCCD970 | $ParA_F$ Q351H-6xHis | pBAD33 | This work |
| pCCD971 | $ParA_F$ W365E-6xHis | pBAD33 | This work |

| | | | |
|---|---|---|---|
| pCCD974 | $ParA_F$ W362E E375A-GFP | pDSW210 | This work |
| pCCD983 | $ParA_F$ W362E R320A-GFP | pDSW210 | This work |
| pCCD892 | $ParA_F$ E375A-GFP | pDSW210 | This work |
| pCCD993 | $ParA_F$ R320A-GFP | pDSW210 | This work |
| pCCD806 | $ParA_F$ E375A-Stop-GFP | pDSW210 | This work |
| pCCD976 | $ParA_F$ W362E E375A-Stop-GFP | pDSW210 | This work |
| pCCD998 | $ParA_F$ R320A-Stop-GFP | pDSW210 | This work |
| pCCD996 | $ParA_F$ W362E R320A-Stop-GFP | pDSW210 | This work |
| pCCD569 | mini-F, cat, $pL_{tetO1}$::*ΔparA, parBS* + | pDAG198 | Castaing et. al., 2008 |

**Supplementary Table S2.**

**List of oligonucleotides used in the study.**

| Primers | Sequence | Description |
|---|---|---|
| RSO425 | 5’ CGGTTCTGGCAAATATTCTGAAATGAGC 3’ | pDSW210-fwd |
| RSO426 | 5’ GCGTTCTGATTTAATCTGTATCAGGC 3’ | pDSW210-rev |
| RSO71 | 5' GTCGTCGTCGACCATGTTCAGAATGAAACTCATG 3' | $ParA_F$ (pREP42) FP |
| RSO72 | 5' GCGGCGTCTAGAGTTGTTGTTGTTTCTAATCTCCCAGCGTGG 3' | $ParA_F$ (pREP42) RP |
| RSO634 | 5’ ATGTTCAGAATGAAACTCATGGAA 3’ | $ParA_F$ FP-Full Length |
| RS0635 | 5’ TCTAATCTCCCAGCGTGGTTTAAT 3’ | $ParA_F$ RP-Full Length |
| RSO753 | 5’ AACAACAACCTGCAGTAAGCTTAAAGGAGAAGAACTTTTCAC 3’ | $ParA_F$-STOP FP |
| RSO754 | 5’ TTGCTTCTCCTTTAAGCTTACTGCAGGTTGTTGTT 3’ | $ParA_F$-STOP RP |

| | | |
|---|---|---|
| RSO98 | 5' CAGTCCCCGTGGATCGAGGAGCAAATTCGGGATGCCTGGGGAAGC 3' | $ParA_F$ M315I fvd |
| RSO99 | 5' CCGAATTTGCTCCTCGATCCACGGGGACTGAGAGCCATTACTATTG 3' | $ParA_F$ M315I Rev |
| RSO460 | 5' CAACTGGTGCCGAGAGAAATGCTCTTTCTATTTGGGAACC 3' | $ParA_F$ W362E fwd |
| RSO461 | 5' GAGCATTTCTCTCGGCACCAGTTGAAGAGCGTTGATCAATGGCC 3' | $ParA_F$ W362E rev |
| RSO100 | 5' CTGTTTTTGAACACGCCATTGATCAACGCTCTTCAACTGGTGCCTGGAG 3' | $ParA_F$ Q351H Fvd |
| RSO101 | 5' CGTTGATCAATGGCGTGTTCAAAAACAGTTCTCATCCGGATC 3' | $ParA_F$ Q351H Rev |
| RSO667 | 5' CCTGTCTGCAATGCAATTTTCGATCGTCTGATT 3' | $ParA_F$ E375A FP |
| RSO668 | 5' ACGATCGAAAATTGCATTGCAGACAGGTTCCCAAATAGA 3' | $ParA_F$ E375A RP |
| RS0990 | 5' GCAAATTgcGGATGCCTGGGGAAGCATG 3' | $ParA_F$ R320A FP |
| RS0991 | 5' GGCATCCgcAATTTGCTCCTCCATCCACGG 3' | $ParA_F$ R320A RP |
| RSO409 | 5' GGTGGCGTTTACGCAACCTCAGTTTCTGTTCATCTTGCTCAGG 3' | $ParA_F$ K120A fwd |
| RSO410 | 5' CAGAAACTGAGGTTGCGTAAACGCCACCTTTATGGGCAGC 3' | $ParA_F$ K120A rev |
| RSO417 | 5' GGTGGCGTTTACCAAACCTCAGTTTCTGTTCATCTTGCTCAGG 3' | $ParA_F$ K120Q_fwd |
| RSO418 | 5' GAAACTGAGGTTTGGTAAACGCCACCTTTATGGGCAGCAAC 3' | $ParA_F$ K120Q_rev |
| RSO863 | 5' GGTGGCGTTTACAGAACCTCAGTTTCTGTTCATCTTGCTCAGG 3' | $ParA_F$ K120R Fwd |
| RSO864 | 5' GAAACTGAGGTTCTGTAAACGCCACCTTTATGGGCAGCAAC 3' | $ParA_F$ K120R Rev |

| | | |
|---|---|---|
| RSO692 | 5' GGTGGCGTTTACGAAACCTCAGTTTCTGTTCATCTTGC TCAGG 3' | $ParA_F$ K120E Fwd |
| RSO693 | 5' GAAACTGAGGTTTCGTAAACGCCACCTTTATGGGCAGC AAC 3' | $ParA_F$ K120E Rev |
